# Signal detectability and boldness are not the same: the function of defensive colouration in nudibranchs is distance-dependent

**DOI:** 10.1101/2022.12.20.521213

**Authors:** Cedric P. van den Berg, John A. Endler, Karen L. Cheney

## Abstract

Aposematic signals advertise underlying defences in many species. They should be detectable (highly contrasting against the background) and bold (high internal pattern contrast) to enhance predator recognition, learning and memorisation. However, the signalling function of aposematic colour patterns may be distance-dependent: signals may be undetectable from a distance to reduce costs of increased attacks from naïve predators but bold when viewed up close. To test this hypothesis, we quantified the chromatic and achromatic detectability and boldness of colour patterns in 13 nudibranch species that varied in the strength of their chemical defences, in terms of unpalatability and toxicity, using Quantitative Colour Pattern Analysis (QCPA) and data on the visual perception of a triggerfish (*Rhinecanthus aculeatus*). When viewed from larger distances, there were no differences in detectability and boldness between well-defended and undefended species. However, when viewed at close distances, well-defended species were more detectable and bolder than undefended species. The detectability of defended species decreased more significantly with increased viewing distances compared to boldness but remained relatively consistent over viewing distances for undefended species. We provide evidence for distance-dependent signalling in aposematic nudibranchs and highlight the importance of distinguishing between signal detectability and boldness in studies of aposematism.

## Introduction

Conspicuous colour patterns displayed by aposematic species are used to educate predators about underlying defences during prey encounters [1,2], or via eavesdropping (e.g. Hämäläinen et al., 2021). Aposematic colour patterns should be readily detectable against their visual background, and detectability is considered crucial to the initial evolution of aposematism [4,5]. Most aposematic signals are also bold, defined as exhibiting high internal colour and luminance contrast [6,7], and striking colour pattern geometry [1,2,8,9], which is thought to enhance the formation and maintenance of predator avoidance behaviour [10]. Theoretically, aposematic colour patterns should be most efficient when they are both easy to detect and are bold [4].

However, it is important to differentiate between detectability and boldness to understand the appearance of a visual signal in the context of its background and the pattern itself. Evidence for or against the relative contribution of each to signal quality and efficacy is mixed [11,12]. For example, Sillén-Tullberg [13] demonstrated that chicks (*Gallus gallus domesticus*) learned to avoid coloured artificial prey independent of background contrast. In contrast, Aronsson and Gamberale-Stille [14] failed to find an effect of internal patterning on the strength of avoidance behaviour in chicks. A key benefit of increased detectability in aposematic species may be to reduce predator recognition errors by allowing more time for accurate decision-making [15]. However, Gamberale-Stille *et al*. [16] demonstrated that increased detectability in chickens (*Gallus gallus domesticus*) did not improve the survival of aposematic prey. In fact, if detectable aposematic species are attacked more frequently by naive predators, or by those that are capable and willing to tolerate secondary defences [e.g. 17,18], detectability may be a potential handicap of bold aposematic signals, rather than a fitness benefit [4,19–23].

To adapt to optimal levels of detectability and signal boldness, aposematic species may benefit from distance-dependent signalling [21,24], which could reduce the detectability of bold colour patterns from a distance while enabling predator deterrence up close. This principle is now supported by multiple theoretical and empirical studies [e.g. 25–31]. For example, Barnett and Cuthill [22] demonstrated that, when searched for by human observers, camouflaged stimuli with boldly contrasting markings were detected at the same distance compared to camouflaged stimuli without bold markings. This was later confirmed by using artificial stimuli and avian predators [32–34].

The apparent paradox of distance-dependent signalling can be investigated by considering the visual perception of a signal in its ecological context, including the spatial acuity of a visual system and the distance at which it is observed [21,35,36]. In the predator sequence, prey detection precedes prey identification and subsequent decisions by predators to further engage with prey [37,38]. Therefore, context-specific signal processing is used by predators as they proceed from detection to discrimination and subsequent attack [12,39,40]. It is likely that the relative importance of primary and secondary defences shifts along such a predation sequence [see 41 for review]. For example, visual defences relying on the avoidance of detection precede bold deimatic displays by threatened prey warning a predator of underlying secondary defences [see 41,42 for reviews]. However, it is unknown how detectability and/or boldness change along an escalating predations sequence in the context of permanently displayed warning signals.

Few studies have differentiated between the detectability and boldness of aposematic animals [but see 43]. Fewer studies have done so at different viewing distances while considering the physiological limitations of ecologically relevant observers [but see 26]. This is partly due to the challenge of capturing spatiochromatic properties of complex visual backgrounds according to observer-specific physiological limitations, such as chromatic and achromatic contrast perception and spatial acuity.

In this study, we investigated the detectability and boldness of defensive animal colouration in 13 species of nudibranch molluscs with differing chemical defences (Fig. 1). Nudibranch molluscs display a stunning diversity of defensive colouration and secondary defences, and are valuable in the study of the ecology and evolution of aposematism [43–46]. We used Quantitative Colour Pattern Analysis (QCPA) [47] and considered the distance at which nudibranchs would be perceived by an ecologically relevant observer, a triggerfish (*Rhinecanthus aculeatus*). We hypothesized that when viewed from larger distances, the detectability of nudibranchs against their backgrounds should be low for all species, irrespective of their bold colour patterns or strength of chemical defences. However, when viewed up close, we hypothesized that species with strong chemical defences would show significantly stronger visual contrasts against their backgrounds than those of undefended species. Lastly, we expected an unequal change in the relationship of boldness and detectability between chemically defended and undefended species across different viewing distances.

**Figure 1:**
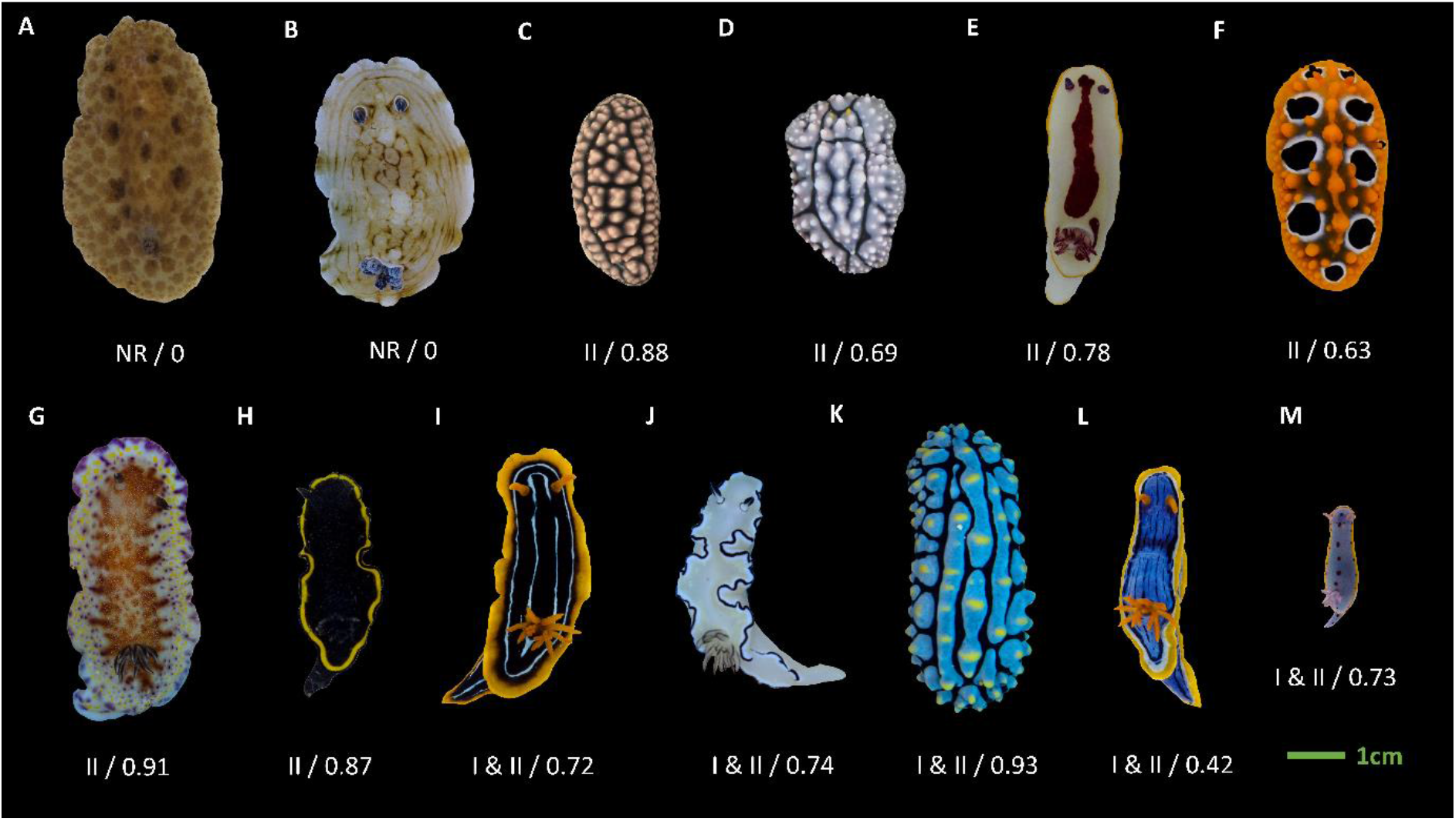
Representative images of species used in this study grouped by their chemical defences scaled approximately according to size: A) *Discodoris sp.; B) Aphelodoris varia; C) Phyllidiella pustulosa; D) Phyllidia elegans; E) Goniobranchus splendidus; F) Phyllidia ocellata; G) Goniobranchus collingwoodi; H) Glossodoris vespa; I) Chromodoris kuiteri; J) Doriprismatica atromarginata; K) Phyllidia varicosa; L) Chromodoris elisabethina; M) Hypselodoris bennetti*. The level of chemical defence is provided as values below each image determined from toxicity and unpalatability assays in Winters et al. 2021. The first value indicates class of chemical defence: ‘NR’ indicates nudibranch species that have limited or no chemical defences; class II’ indicate species that are highly unpalatable but weakly toxic and class ‘I & II’ indicate species are highly unpalatable and highly toxic. The second value is data from unpalatability feeding assays with Palaemon shrimp (presented here as 1-ED_50_ values). ED_50_ refers to the median effective dose.

## 2. Material and Methods

### (a) Image collection

We took calibrated digital images of 13 Dorid nudibranch species (n = 226 individuals): *Aphelodoris varia* (n = 24), *Chromodoris elisabethina* (n = 21), *Chromodoris kuiteri* (n = 17), *Discodoris sp*. (n = 15), *Doriprismatica atromarginata* (n = 27), *Glossodoris vespa* (n = 15), *Goniobranchus collingwoodi* (n = 15), *Goniobranchus splendidus* (n = 25), *Hypselodoris bennetti* (n = 10), *Phyllidia elegans* (n = 8)*, Phyllidia ocellata* (n = 23), *Phyllidia varicosa* (n = 8) and *Phyllidiella pustulosa* (n = 18). Species were visually identified using various taxonomic books [48–50]. We grouped individuals that visually resembled *Sebadoris fragilis* and were found in the same locations as *Discodoris* sp.; however, these individuals could be a mixture of *Sebadoris fragilis, Thayuva lilacina, Jorunna pantheris* and perhaps other undescribed species that cannot be identified without molecular sequencing.

Fieldwork was conducted on SCUBA between October 2017 and July 2021 at dive sites along the east coast of Australia: Sunshine Coast (SE Queensland), Gold Coast (SE Queensland) and Nelson Bay (New South Wales). Individual nudibranchs were located and photographed underwater at depths of 2-18m against their natural habitat using a calibrated digital Olympus EPL-5 with a 60mm macro lens in an Olympus PT-EP10 underwater housing. We used white LED illumination from a combination of VK6r and PV62 Scubalamp video lights for illumination [47]. All images were taken at roughly a 90° angle (top-down) relative to the animal and its background with the animals in a naturally stretched-out, straight, forward-moving position (Fig. 1).

### (b) Aposematism and chemical defences

With the exception of *Discodoris sp*. and *Aphelodoris varia*, we consider these species to be aposematic as they harbour potent chemical defences and display bold colour signals [46,51] Fig. 1). *Aphelodoris varia* has no known chemical defences [46] and likely relies on crypsis via background matching for their primary defence. *Sebadoris fragilis* does not have any known chemical defences [46] and we, therefore, treat *Discodoris sp*. as non-toxic and palatable.

We quantified chemical defences in two ways in our study. First, we used the classification from [46] to delineate between species belonging to the following three classes: NR - nudibranch species that have limited or no chemical defences, class ‘II’ – species that are highly unpalatable but only weakly toxic, class ‘I & II’ – highly unpalatable and highly toxic. *Glossodoris vespa* was classified as class II from assay data reported in [51]. Second, we ranked the species from 0 to 1, according to the average ED_50_ response in *Palaemon* shrimp feeding assays reported in [46,51], calculated as 1-ED_50_ so that 1 represented the most unpalatable species (Fig 1).

### (b) Quantification of detectability and boldness

Image analysis was conducted with visual modelling parameter choices as per [52] at the following viewing distances: 2cm, 5cm, 10cm and 30cm. At distances beyond 30cm, the majority of small nudibranchs and much of the internal patterning in larger individuals are unlikely to be visible to a triggerfish (*R. aculeatus*) due to its spatial acuity of about 3 cycles per degree (cpd) [53].

To quantify detectability (i.e. a measure of background matching), we calculated the absolute difference of the abundance weighted coefficient of variation of achromatic (Lum.CoV) and chromatic (Col.CoV) local edge contrast (LEIA, measured in ΔS) for each animal relative to that of its respective visual background. Thus, by considering the difference in the overall appearance of an animal by itself relative to its immediate background, we specifically quantify background matching. The coefficient of variation of chromatic and achromatic boundary contrast between colour pattern elements has previously been shown to indicate mate choices in guppies (*Poecilia reticulata*) [54,55]. Instead of using the boundary strength analysis [BSA, 56], we make use of the non-parametric approach of LEIA and its ability to consider colour pattern contrast at roughly the scale of an edge-detecting receptive field of a triggerfish [57]. Unlike BSA, LEIA does not rely on image segmentation. After applying Gaussian acuity modelling, we use receptor noise limited (RNL) ranked filtering [47] as per [52].

The RNL model [58] allows us to estimate the perceived colour and luminance contrast of non-human observers and, in addition to RNL ranked filtering, is used to describe the contrast of edges within each animal and its corresponding natural background. This contrast measure is expressed as ΔS and reflects the distance between two contrasting points in the RNL colour space [59]. To describe the colour and luminance contrast at each location in the image, we used the average LEIA contrast across the vertical, horizontal and diagonal filter axis. For a detailed description of LEIA parameters, see [47].

To quantify boldness, we used Lum.Cov and Col.CoV of each animal without considering their respective visual backgrounds, as we define the boldness of an aposematic signal as the strength of the signal once prey detection has taken place. High values for each of these pattern statistics indicate highly contrasting colour pattern elements.

### (c) Statistical analysis

All colour pattern statistics were normalised to range from 0-1 using the ‘range’ argument of the PreProcess function (caret package, [60], v6.0-90) using R Software ([61], v4.1.2), with infinite and NA values treated as 0.

When we considered differences between species without considering the class of their chemical defences, detectability and boldness data did not meet the assumptions of normality due to large variation in the number of individuals and individual trait data, so we used a Kruskal-Wallis test [62]. This analysis was performed using the *stats* package and post hoc analyses comparing individuals was done using Dunn tests [63] with a Bonferroni correction [64] in the *FSA* package ([65], v0.9.3).

When considering differences between species in relation to their class of chemical defence at different viewing distances, we applied a linear mixed effect (lme) model using the lme4 [66], v1.1-28) and lmertest [67], v3.1-3) packages and type III ANOVA tables using Saterwaite’s method in the stats package after applying a square-root transform to the left-skewed data. The meeting of assumptions was assessed for each model fit. We further ran independent lme models for chemical defence and viewing distance. Species was treated as a random effect for all lme models.

To determine the relationship between unpalatability data and colour pattern statistics, we used a Pearson product-moment correlation (R) in the stats package [61], v4.1.2). Detectability and boldness were measured as the species median value of Lum.Cov and Col.Cov at each viewing distance. For this analysis, we applied an ordered quantile normalisation to the raw data using the bestNormalize package [68], v.1.8.2) to ensure normality throughout the dataset. The distance-dependent change in the correlation between unpalatability and visual defences was then assessed by fitting linear regressions to the obtained Pearson product-moment correlations.

## 3. Results

### (a) Detectability

We first considered differences in detectability between species (i.e. the difference between each animal and its background) at different distances without considering chemical defences. We found significant differences in achromatic detectability (Lum.CoV) between species at all viewing distances (Kruskal-Wallis 2cm: *χ*^2^_(12)_ = 126.35, *p* < 0.001; 5cm: *χ*^2^_(12)_ = 111.44, *p* < 0.001; 10cm: *χ*^2^_(12)_ = 75.389, *p* < 0.001; 30cm: *χ*^2^_(12)_ = 24.94, *p* = 0.015). However, only one species remained significantly more detectable than others at 30cm, with *H. bennetti* being more detectable than *G. vespa* (Dunn-test: *p_adj_* = 0.021, Table S1). Similarly, there were significant differences in chromatic detectability between species at close viewing distances (Kruskal-Wallis 2cm: *χ*^2^_(12)_ = 95.23, *p* < 0.001; 5cm: *χ*^2^_(12)_ = 97.69, *p* < 0.001; 10cm: *χ*^2^_(12)_ = 76.81, *p* < 0.001) but not at 30cm (Kruskal-Wallis: *χ*^2^_(12)_ = 16.62, *p* = 0.164, Table S2)

When considering the class of chemical defence, species without chemical defences (NR) had higher achromatic detectability (Lum.Cov) at larger viewing distances (F_4,189_ = 6.27, *p* < 0.001). In contrast, chemically defended species were less detectable when viewed from a distance, with the most defended species (class I and II) displaying the largest differences between viewing distances (weakly toxic / highly unpalatable (class II): *F_4,510_* = 12.31, *p* < 0.001; highly toxic / highly unpalatable (class I & II): *F_4,406_* = 31.38, *p* < 0.001) (Fig. 2a).

**Figure 2.**
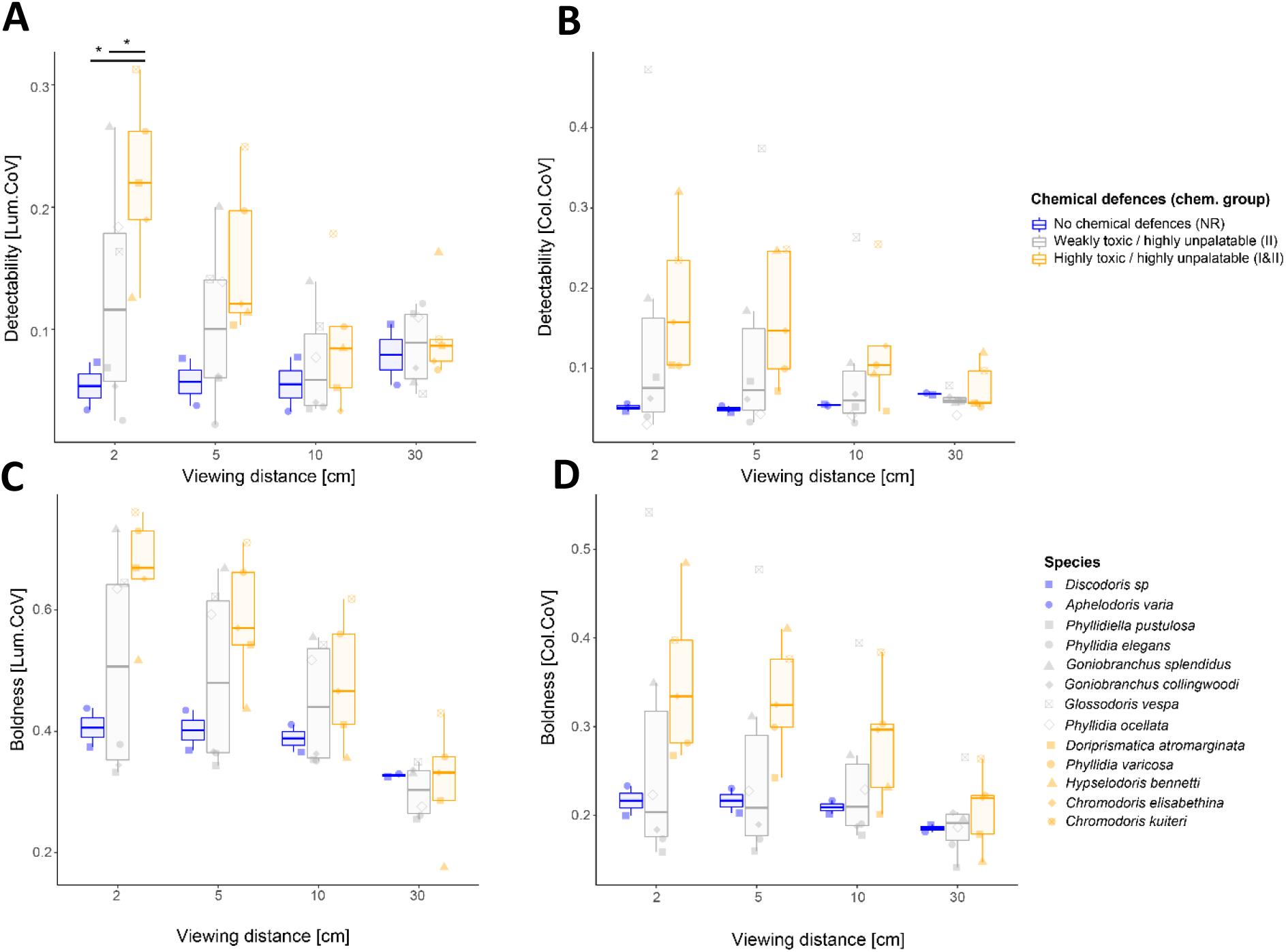
Relative detectability and boldness of 13 Dorid nudibranch species. The strength of chemical defences is indicated by colours relative to the class of chemical defence. A) Species median detectability quantified as the abundance weighted coefficient of variation of achromatic local edge contrast (Lum.CoV) measured relative to the natural background of every individual. B) Species median detectability quantified as the abundance weighted coefficient of variation of chromatic local edge contrast (Col.CoV) measured relative to the natural background of every individual. C) Species median achromatic pattern boldness measured as the abundance weighted coefficient of variation of achromatic local edge contrast (Lum.CoV) of each animal. D) Species median chromatic pattern boldness measured as the as the abundance weighted coefficient of variation of chromatic local edge contrast (Col.CoV) of each animal.

For species without chemical defences (NR), there was no difference in chromatic detectability (Col.Cov) between viewing distances (*F_4,190_* = 0.50, *p* = 0.734). In contrast, species with chemical defences (classes II / I & II) showed a strong reduction in chromatic detectability with increasing viewing distances (weakly toxic / highly unpalatable (class II): *F_4,510_* = 12.24, *p* < 0.001; highly toxic / highly unpalatable (class I & II): *F_4,509.98_* = 16.13, *p* < 0.001). Again, this relationship was stronger for species with both toxic and unpalatable chemical defences (class I & II) (Fig. 2b).

There was no significant differences in either achromatic (Lum.CoV - 5cm: *F_2,9.94_* = 2.07, *p* = 0.178, 10cm: *F_2,10.17_* = 0.27, *p* = 0.771, 30cm: *F_2,8.58_* = 0.07, *p* = 0.937) or chromatic detectability (Col.CoV - 2cm: *F_2,9.82_* = 1.09, *p* = 0.374, 5cm: *F_2,9.88_* = 1.03, *p* = 0.391, 10cm: *F_2,9.95_* = 0.75, *p* = 0.498, 30cm: *F_2,8.68_* = 0.22, *p* = 0.807) between chemical defence groups at either viewing distance. However, at 2cm achromatic detectability species with both toxic and unpalatable chemical defences (class I & II) were more detectable than either species without chemical defences (class NR) or unpalatable ones (class II) (*F_2,9.8_* = 4.26, *p* = 0.047).

Unpalatability data was weakly positively correlated with both achromatic and chromatic detectability at close viewing distances, whereas no or weakly negative correlations were observed at larger viewing distances (Fig. 3a). Both, achromatic (*r^2^*= 0.98, *F_1,2_*= 117.7, *p* = 0.008) and chromatic (*r^2^*= 0.98, *F_1,2_*= 110, *p* = 0.009) detectability showed a significant reduction in their correlation with unpalatability over increasing viewing distances (Fig. 3b).

**Figure 3.**
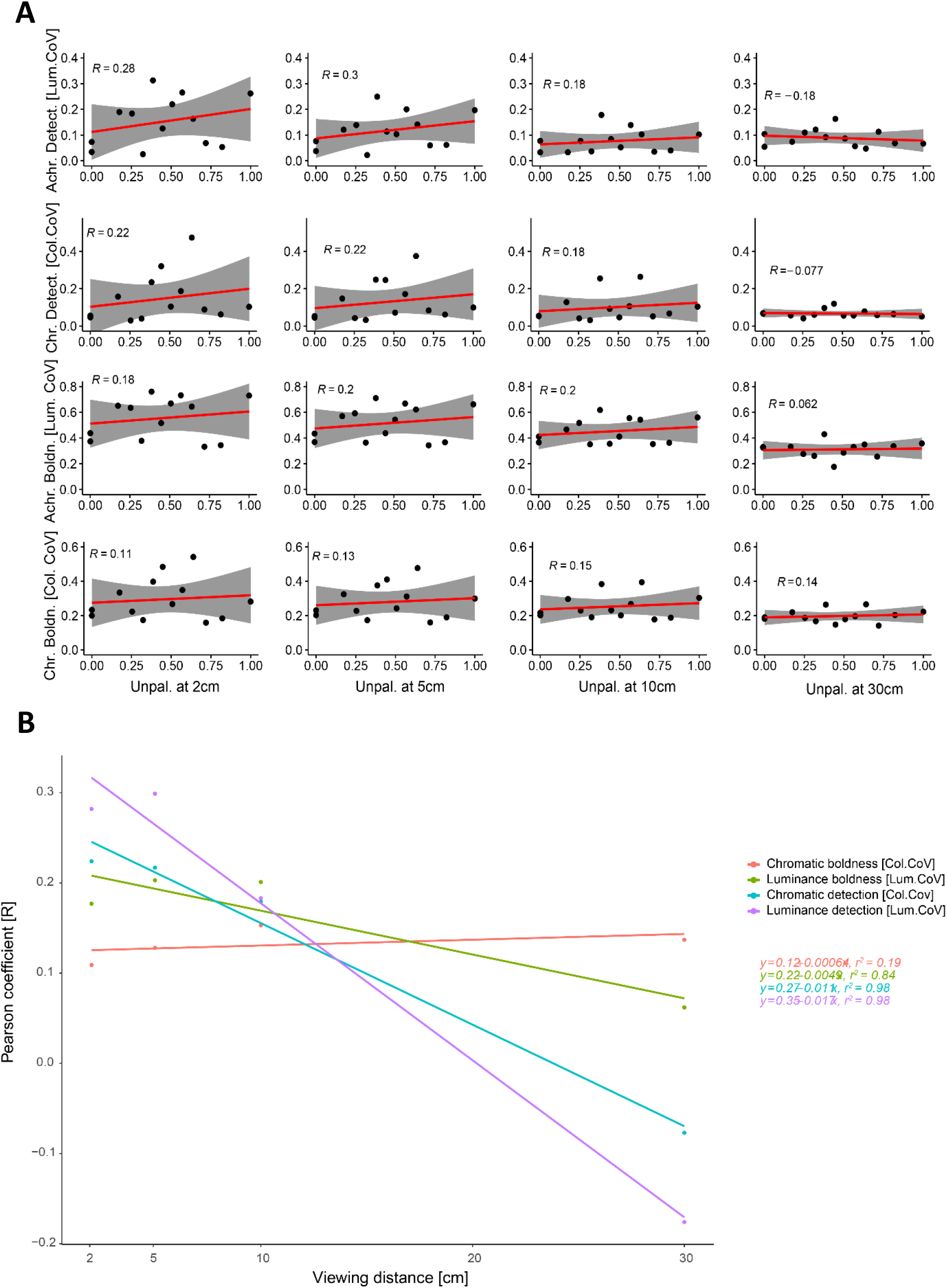
Summary of the correlation of the median chromatic (Col.CoV) and achromatic (Lum.CoV) boldness and detectability of each species with the strength of chemical defences (1-ED_50_) across different viewing distances. A) Individual Pearson product-moment correlations (R) for each viewing distance and colour pattern statistic. Grey shading shows 95% confidence intervals; B) Linear regressions fitted to the Pearson product-moment correlations (R) show variable distance-dependent decreases in the correlation between colour pattern statistics.

### (b) Boldness

When we considered differences in boldness (i.e. analysing each animal without its background) without considering chemical defences, we found significant differences in achromatic boldness between species at all viewing distances (Kruskal-Wallis 2cm:*χ*^2^_(12)_ = 159.94, *p* < 0.001; 5cm: *χ*^2^_(12)_ = 161.11, *p* < 0.001; 10cm: *χ*^2^_(12)_ = 148.26, *p* < 0.001; 30cm: *χ*^2^_(12)_ = 74.27, *p* < 0.001). At 30cm, significant differences between multiple species remained (Fig. 2c, Table S3). Similarly, there were significant differences in chromatic boldness between species across all viewing distances (Kruskal-Wallis 2cm: *χ*^2^_(12)_ = 165.46, *p* < 0.001; 5cm: *χ*^2^_(12)_ = 163.55, *p* < 0.001; 10cm: *χ*^2^_(12)_ = 147.8, *p* < 0.001, 30cm: *χ*^2^_(12)_ = 88.89, *p* < 0.001) (Table S4).

When considering the class of chemical defence, species with and without chemical defences showed decreased achromatic boldness at larger viewing distances, with the most defended species displaying the largest differences between distances (Undefended (class NR): *F_4,189_* = 38.34, *p* < 0.001; weakly toxic / highly unpalatable (class II): *F_4,510.05_* = 244.74, *p* < 0.001; highly toxic / highly unpalatable (class I & II): *F_4,406_* = 324.83, *p* < 0.001) (Fig. 2).

Similarly, species with and without chemical defences showed decreased chromatic boldness at larger viewing distances with the most defended species displaying the largest differences between distances (Undefended (NR): *F_4,189_* = 6.27, *p* < 0.001; Weakly toxic / highly unpalatable (II): *F_4,510_* = 12.31, *p* < 0.001; Highly toxic / highly unpalatable (I & II): *F_4,406_* = 31.38, *p* < 0.001) (Fig. 2d).

There were no significant differences in either achromatic (Lum.CoV - 2cm: *F_2,9.95_* = 2.43, *p* = 0.139, 5cm: *F_2,9.92_* = 1.33, *p* = 0.309, 10cm: *F_2,9.87_* = 0.34, *p* = 0.721, 30cm: *F_2,9.48_* = 0.19, *p* = 0.828) or chromatic boldness (Col.CoV - 2cm: *F_2,9.95_* = 1.08, *p* = 0.375, 5cm: *F_2,9.97_* = 1.28, *p* = 0.320, 10cm: *F_2,9.80_* = 0.70, *p* = 0.519, 30cm: *F_2,9.77_* = 0.05, *p* = 0.947) between chemical defence groups at either viewing distance.

When considering chemical defences on a continuous scale using unpalatability data, achromatic boldness showed a moderate correlation with unpalatability at close viewing distances whereas chromatic boldness showed a weak correlation at close viewing distances (Fig. 3a). However, neither achromatic (*r^2^*= 0.84, *F_1,2_*= 10.38, *p* = 0.084) nor chromatic (*r^2^*= 0.19, *F_1,2_*= 0.48, *p* = 0.560) detectability showed a significant reduction in their correlation with unpalatability over increasing viewing distances (Fig. 3b).

## 4. Discussion

Our study demonstrates that the appearance of nudibranch colouration in the context of natural backgrounds (detectability) or by itself (boldness) depends on the distance at which it is viewed by an animal observer. For highly defended nudibranchs, both achromatic and chromatic detectability and boldness significantly reduced with increasing viewing distance. In contrast, the detectability and boldness of undefended species remained relatively consistent over viewing distances. Our results support predictions [21,24] and empirical findings [e.g. 22] suggesting that colour patterns with bold markings can be as camouflaged at a distance as colour patterns that are camouflaged using background matching.

While chromatic and achromatic boldness decreased equally with increasing viewing distances, various significant differences in achromatic and chromatic boldness remained between species when viewed at 30 cm (Fig. 2). This could indicate that some species remain distinct in appearance at viewing distances where the likelihood of detection is already significantly reduced. Therefore, some well-defended species might still profit from a bold appearance in the unlikely event of being detected at larger distances. These findings align with the suggestion from Barnett *et al*. [34] that detectability and signalling function of defensive colouration can be differentially influenced by viewing distance and that a reduction in detectability does not necessarily mean an equal reduction in signal quality. Our study is the first to support this hypothesis using animals observed in their natural habitat.

The loss of perceived higher spatial frequencies with increasing viewing distance affects both apparent features and detail in visual backgrounds, as well as within an animal. As viewing distances increase, smaller-scale colour pattern elements are expected to blur together, leading to the additive blending of colour pattern elements [69] and subsequent change in the visual appearance of colour patterns and visual backgrounds. How, and if, such blending happens, is influenced by the size, shape and colour of individual colour pattern elements. However, in many animals chromatic detail is perceived at lower spatial acuity than achromatic contrast [e.g. 70]. Thus, how chromatic and achromatic contrast is arranged in colour patterns can allow for independently variable detectability or boldness to a given predator at a given distance. Furthermore, object detection and discrimination rely on vastly different processing pathways with object detection not necessarily requiring object recognition [71]. As a result, individual or multiple colour pattern design elements can be subject to individual or multiple selective pressures [39,72]. Therefore, despite being influenced by the loss of spatial detail as viewing distances increase, colour pattern boldness and detectability in aposematic animals may not be subject to equal selective pressures at varying viewing distances [e.g. 34].

Regulating the relative importance of detectability and boldness has been proposed to be crucial in aposematic species to balance the costs and benefits of increased detectability induced by bold colouration [34]. Therefore, detectability and boldness are thought to coincide with aposematism at close viewing distances whereas aposematic and cryptic species are likely to profit from camouflage at a distance [21,24]. Our findings support this assumption, as animals with both toxic and unpalatable chemical defences were more detectable at close viewing distances than animals lacking chemical defences or those with only unpalatable chemical defences (Fig. 3). We also did not find significant differences in detectability between species with chemical defences and no chemical defences at larger viewing distances (Fig. 3). However, while there were limited differences in detectability between species at 30cm viewing distance, substantial variation in terms of chromatic and achromatic boldness remained between species (Fig. 2, Table S3 & S4). Our findings suggest that the boldness and detectability of colour patterns in aposematic nudibranch molluscs show distinct distance-dependent correlations with increasing viewing distances. In line with predictions from Barnett et al. (2017), our results show that detectability in chemically defended nudibranchs decreases faster than boldness as viewing distances increase (Fig. 3). Specifically, both chromatic and achromatic detectability show a significant decrease in their correlation with unpalatability, whereas boldness does not. This highlights the cumulative importance of distinguishing between both, signal detectability and boldness as well as ecologically relevant viewing distances when investigating signalling honesty in aposematic species. However, it is important to note that the relative importance of detectability and colour pattern boldness is context specific, as a bold animal is not necessarily easy to detect in a visual background containing similar levels of colour pattern contrast [21,73,74], such is the case, for example, if the animal’s colour pattern represents a random sample of its complex and highly contrasty visual background [21].

Interestingly, the intersection of the distance-dependent decreases of the correlation between detectability and boldness is situated at around 10cm (Fig. 3). At this distance, many of the more intricate colour pattern details are no longer visible to triggerfish. This likely hinders the identification of and discrimination between species by potential predators, further emphasising the importance of distinguishing between cognitive processes involved in prey detection and identification. Such distance-dependent effects on the perception of colour patterns could be crucial in the ecology and evolution of mimicry [75]. For example, the bold, outline-enhancing yellow rims displayed by some of the species in this study are likely to be more detectable at a distance than the finer internal patterning. This could explain the presence and constancy of such outline enhancing yellow rims [9] among many aposematic nudibranch species and their putative mimics [72].

We show that the distance-dependent correlation between boldness and detectability depends on the definition and strength of both underlying chemical and visual defences (Fig. 2 & 3). For example, when correlating unpalatability on a continuous scale with each species’ median colour pattern statistic, we found that the correlation between detectability and the strength of chemical defences decreases significantly with increasing viewing distances, whereas this does not happen for boldness (Fig. 3). Instead, the correlation between boldness and unpalatability stays constant as viewing distance increases. However, when looking at chemical defences defined as categories considering both unpalatability and toxicity, we find significant decreases in boldness for chemically defended species (Fig. 2). Furthermore, the effect size for the decrease of achromatic boldness is much larger than that for the decrease of chromatic boldness. Therefore, in addition to highlighting the importance of considering achromatic and chromatic aspects of aposematic signals separately, our results highlight the importance of considering the type of analysis applied to such data.

As with many aposematic animals [see 76 for review], colour patterns within species of Dorid nudibranchs are known to vary substantially [e.g. 72,77]. Furthermore, species of Dorid nudibranchs display substantial degrees of seasonal and geographic variation in their chemical defences [46,72] and sample sizes (n = 1-3 extractions of pooled individuals per species) in [46] are unlikely to capture this variability. However, larger sample sizes for each species for both, chemical defences and colour patterns, will be of great interest in future studies. Despite such limitations, our results show that considering individual variability as opposed to using species averages and defining chemical defences in terms of categories as opposed to a continuous scale shapes the conclusions on the distance-dependent signalling function of aposematic animals. This is especially true considering that, despite considering many of the most common species, this study only captures 13 out of hundreds of species of sea slugs present in the study area [78]

Perceptual mechanisms underlying prey detection, identification and assessment are complex and remain insufficiently quantified in studies of visual ecology [41,47,52,79]. Furthermore, rather than considering the entire colour pattern space available in QCPA we quantify detectability and boldness using a pre-determined selection of image statistics based on existing literature. These statistics are generated using a specific set of modelling parameters guided by the best available knowledge. However, a significant amount of work remains to be done to identify relevant visual features and modelling parameters in each behavioural context [52,79]. Therefore, revisiting distance dependent signalling in nudibranch molluscs using a larger dataset with a more informed approach on modelling and statistical analysis will remain of great interest for future research. And yet, our study and represents the currently most common approach towards the quantification of visual signals in our field, even when using analyses such as QCPA [e.g. 80,81].

In summary, our results suggest that many aposematic Dorid nudibranchs are equally as cryptic by means of background matching as undefended species at greater viewing distances. Our results further suggest a multifunctional, multicomponent design of colour patterns in aposematic Dorid nudibranchs. Moreover, our findings emphasize the need for a cautious and well-defined use of the term conspicuousness when discussing aposematism. It is important to distinguish between the detectability and boldness of aposematic signals when investigating colour pattern functionality, particularly when investigating signal honesty in aposematic species [82,83]. However, differential selection on detectability and colour pattern boldness likely also applies to other forms of visual signalling in prey animals, such as sexual or territorial signalling, providing a great area of investigation for future studies.

## Data accessibility

The data will be uploaded to UQ’s e-space upon acceptance.

## Author’s contributions

CPvdB conceived the original study with KLC & JAE providing critical input; CPvdB collected and analysed the data; CPvdB and KLC led the writing of the manuscript. All authors contributed critically to the drafts and gave final approval for publication.

## Acknowledgements

We would like to thank Dr Nicholas Condon for his assistance with the creation of automated QCPA scripts, various volunteers for assistance with image analysis and fieldwork. We also thank the staff running the High-Performance Computing infrastructure at The University of Queensland (Wiener & Awoonga) and Dr. Simone Blomberg, who provided infrastructure that contributed to the computing of image statistics.

