## Supplement for "Signal detectability and boldness are not the same: the function of defensive colouration in nudibranchs is distance-dependent"

**Table S1.** Dunn tests at species level for achromatic detectability (Lum.CoV). Significant (Bonferroni adjusted p) pairwise comparisons in red.

| **2cm** | | | | | **5cm** | | | | | **10cm** | | | | | **30cm** | | | |
| --- | --- | --- | --- | --- | --- | --- | --- | --- | --- | --- | --- | --- | --- | --- | --- | --- | --- | --- |
| Comparison | Z | P.unadj | P.adj | Comparison | | Z | P.unadj | P.adj | Comparison | | Z | P.unadj | P.adj | Comparison | | Z | P.unadj | P.adj |
| Aphelodoris varia - Chromodoris elisabethina | -5.069 | <0.001 | <0.001 | Aphelodoris varia - Chromodoris elisabethina | | -3.792 | <0.001 | 0.012 | Aphelodoris varia - Chromodoris elisabethina | | -0.689 | 0.491 | 1 | Aphelodoris varia - Chromodoris elisabethina | | -0.526 | 0.599 | 1 |
| Aphelodoris varia - Chromodoris kuiteri | -7.276 | <0.001 | <0.001 | Aphelodoris varia - Chromodoris kuiteri | | -7.2 | <0.001 | 0 | Aphelodoris varia - Chromodoris kuiteri | | -5.893 | <0.001 | <0.001 | Aphelodoris varia - Chromodoris kuiteri | | -0.114 | 0.909 | 1 |
| Chromodoris elisabethina - Chromodoris kuiteri | -2.427 | 0.015 | 1 | Chromodoris elisabethina - Chromodoris kuiteri | | -3.523 | <0.001 | 0.033 | Chromodoris elisabethina - Chromodoris kuiteri | | -5.095 | <0.001 | <0.001 | Chromodoris elisabethina - Chromodoris kuiteri | | 0.371 | 0.711 | 1 |
| Aphelodoris varia - Discodoris sp | -1.385 | 0.166 | 1 | Aphelodoris varia - Discodoris sp | | -2.093 | 0.036 | 1 | Aphelodoris varia - Discodoris sp | | -2.889 | 0.004 | 0.302 | Aphelodoris varia - Discodoris sp | | -1.577 | 0.115 | 1 |
| Chromodoris elisabethina - Discodoris sp | 3.132 | 0.002 | 0.136 | Chromodoris elisabethina - Discodoris sp | | 1.314 | 0.189 | 1 | Chromodoris elisabethina - Discodoris sp | | -2.204 | 0.028 | 1 | Chromodoris elisabethina - Discodoris sp | | -1.07 | 0.285 | 1 |
| Chromodoris kuiteri - Discodoris sp | 5.224 | <0.001 | <0.001 | Chromodoris kuiteri - Discodoris sp | | 4.499 | <0.001 | 0.001 | Chromodoris kuiteri - Discodoris sp | | 2.59 | 0.01 | 0.75 | Chromodoris kuiteri - Discodoris sp | | -1.363 | 0.173 | 1 |
| Aphelodoris varia - Doriprismatica atromarginata | -6.3 | <0.001 | <0.001 | Aphelodoris varia - Doriprismatica atromarginata | | -3.95 | <0.001 | 0.006 | Aphelodoris varia - Doriprismatica atromarginata | | -1.309 | 0.19 | 1 | Aphelodoris varia - Doriprismatica atromarginata | | -1.068 | 0.285 | 1 |
| Chromodoris elisabethina - Doriprismatica atromarginata | -0.869 | 0.385 | 1 | Chromodoris elisabethina - Doriprismatica atromarginata | | 0.086 | 0.931 | 1 | Chromodoris elisabethina - Doriprismatica atromarginata | | -0.555 | 0.579 | 1 | Chromodoris elisabethina - Doriprismatica atromarginata | | -0.489 | 0.625 | 1 |
| Chromodoris kuiteri - Doriprismatica atromarginata | 1.741 | 0.082 | 1 | Chromodoris kuiteri - Doriprismatica atromarginata | | 3.793 | <0.001 | 0.012 | Chromodoris kuiteri - Doriprismatica atromarginata | | 4.847 | <0.001 | <0.001 | Chromodoris kuiteri - Doriprismatica atromarginata | | -0.851 | 0.395 | 1 |
| Discodoris sp - Doriprismatica atromarginata | -4.072 | <0.001 | 0.004 | Discodoris sp - Doriprismatica atromarginata | | -1.302 | 0.193 | 1 | Discodoris sp - Doriprismatica atromarginata | | 1.812 | 0.07 | 1 | Discodoris sp - Doriprismatica atromarginata | | 0.681 | 0.496 | 1 |
| Aphelodoris varia - Glossodoris vespa | -3.882 | <0.001 | 0.008 | Aphelodoris varia - Glossodoris vespa | | -4.203 | <0.001 | 0.002 | Aphelodoris varia - Glossodoris vespa | | -3.679 | <0.001 | 0.018 | Aphelodoris varia - Glossodoris vespa | | 0.688 | 0.492 | 1 |
| Chromodoris elisabethina - Glossodoris vespa | 0.701 | 0.484 | 1 | Chromodoris elisabethina - Glossodoris vespa | | -0.74 | 0.459 | 1 | Chromodoris elisabethina - Glossodoris vespa | | -2.973 | 0.003 | 0.23 | Chromodoris elisabethina - Glossodoris vespa | | 1.135 | 0.256 | 1 |
| Chromodoris kuiteri - Glossodoris vespa | 2.904 | 0.004 | 0.288 | Chromodoris kuiteri - Glossodoris vespa | | 2.538 | 0.011 | 0.869 | Chromodoris kuiteri - Glossodoris vespa | | 1.856 | 0.064 | 1 | Chromodoris kuiteri - Glossodoris vespa | | 0.741 | 0.458 | 1 |
| Discodoris sp - Glossodoris vespa | -2.251 | 0.024 | 1 | Discodoris sp - Glossodoris vespa | | -1.902 | 0.057 | 1 | Discodoris sp - Glossodoris vespa | | -0.712 | 0.476 | 1 | Discodoris sp - Glossodoris vespa | | 2.041 | 0.041 | 1 |
| Doriprismatica atromarginata - Glossodoris vespa | 1.52 | 0.128 | 1 | Doriprismatica atromarginata - Glossodoris vespa | | -0.855 | 0.393 | 1 | Doriprismatica atromarginata - Glossodoris vespa | | -2.619 | 0.009 | 0.688 | Doriprismatica atromarginata - Glossodoris vespa | | 1.633 | 0.102 | 1 |
| Aphelodoris varia - Goniobranchus collingwoodi | -1.358 | 0.175 | 1 | Aphelodoris varia - Goniobranchus collingwoodi | | -1.033 | 0.301 | 1 | Aphelodoris varia - Goniobranchus collingwoodi | | -1.513 | 0.13 | 1 | Aphelodoris varia - Goniobranchus collingwoodi | | -0.115 | 0.909 | 1 |
| Chromodoris elisabethina - Goniobranchus collingwoodi | 3.159 | 0.002 | 0.124 | Chromodoris elisabethina - Goniobranchus collingwoodi | | 2.346 | 0.019 | 1 | Chromodoris elisabethina - Goniobranchus collingwoodi | | -0.865 | 0.387 | 1 | Chromodoris elisabethina - Goniobranchus collingwoodi | | 0.354 | 0.724 | 1 |
| Chromodoris kuiteri - Goniobranchus collingwoodi | 5.25 | <0.001 | <0.001 | Chromodoris kuiteri - Goniobranchus collingwoodi | | 5.483 | <0.001 | <0.001 | Chromodoris kuiteri - Goniobranchus collingwoodi | | 3.868 | <0.001 | 0.009 | Chromodoris kuiteri - Goniobranchus collingwoodi | | -0.004 | 0.997 | 1 |
| Discodoris sp - Goniobranchus collingwoodi | 0.025 | 0.98 | 1 | Discodoris sp - Goniobranchus collingwoodi | | 0.955 | 0.34 | 1 | Discodoris sp - Goniobranchus collingwoodi | | 1.24 | 0.215 | 1 | Discodoris sp - Goniobranchus collingwoodi | | 1.318 | 0.188 | 1 |
| Doriprismatica atromarginata - Goniobranchus collingwoodi | 4.101 | <0.001 | 0.003 | Doriprismatica atromarginata - Goniobranchus collingwoodi | | 2.384 | 0.017 | 1 | Doriprismatica atromarginata - Goniobranchus collingwoodi | | -0.406 | 0.685 | 1 | Doriprismatica atromarginata - Goniobranchus collingwoodi | | 0.813 | 0.416 | 1 |
| Glossodoris vespa - Goniobranchus collingwoodi | 2.276 | 0.023 | 1 | Glossodoris vespa - Goniobranchus collingwoodi | | 2.857 | 0.004 | 0.334 | Glossodoris vespa - Goniobranchus collingwoodi | | 1.952 | 0.051 | 1 | Glossodoris vespa - Goniobranchus collingwoodi | | -0.723 | 0.47 | 1 |
| Aphelodoris varia - Goniobranchus splendidus | -6.942 | <0.001 | <0.001 | Aphelodoris varia - Goniobranchus splendidus | | -7.07 | <0.001 | <0.001 | Aphelodoris varia - Goniobranchus splendidus | | -5.351 | <0.001 | <0.001 | Aphelodoris varia - Goniobranchus splendidus | | -1.947 | 0.051 | 1 |
| Chromodoris elisabethina - Goniobranchus splendidus | -1.585 | 0.113 | 1 | Chromodoris elisabethina - Goniobranchus splendidus | | -2.998 | 0.003 | 0.212 | Chromodoris elisabethina - Goniobranchus splendidus | | -4.471 | <0.001 | 0.001 | Chromodoris elisabethina - Goniobranchus splendidus | | -1.349 | 0.177 | 1 |
| Chromodoris kuiteri - Goniobranchus splendidus | 1.026 | 0.305 | 1 | Chromodoris kuiteri - Goniobranchus splendidus | | 0.833 | 0.405 | 1 | Chromodoris kuiteri - Goniobranchus splendidus | | 1.078 | 0.281 | 1 | Chromodoris kuiteri - Goniobranchus splendidus | | -1.655 | 0.098 | 1 |
| Discodoris sp - Goniobranchus splendidus | -4.678 | 0 | 0 | Discodoris sp - Goniobranchus splendidus | | -4.077 | <0.001 | 0.004 | Discodoris sp - Goniobranchus splendidus | | -1.771 | 0.077 | 1 | Discodoris sp - Goniobranchus splendidus | | -0.115 | 0.909 | 1 |
| Doriprismatica atromarginata - Goniobranchus splendidus | -0.779 | 0.436 | 1 | Doriprismatica atromarginata - Goniobranchus splendidus | | -3.287 | 0.001 | 0.079 | Doriprismatica atromarginata - Goniobranchus splendidus | | -4.186 | <0.001 | 0.002 | Doriprismatica atromarginata - Goniobranchus splendidus | | -0.925 | 0.355 | 1 |
| Glossodoris vespa - Goniobranchus splendidus | -2.162 | 0.031 | 1 | Glossodoris vespa - Goniobranchus splendidus | | -1.951 | 0.051 | 1 | Glossodoris vespa - Goniobranchus splendidus | | -0.975 | 0.33 | 1 | Glossodoris vespa - Goniobranchus splendidus | | -2.397 | 0.017 | 1 |
| Goniobranchus collingwoodi - Goniobranchus splendidus | -4.706 | <0.001 | <0.001 | Goniobranchus collingwoodi - Goniobranchus splendidus | | -5.145 | <0.001 | <0.001 | Goniobranchus collingwoodi - Goniobranchus splendidus | | -3.157 | 0.002 | 0.124 | Goniobranchus collingwoodi - Goniobranchus splendidus | | -1.588 | 0.112 | 1 |
| Aphelodoris varia - Hypselodoris bennetti | -2.452 | 0.014 | 1 | Aphelodoris varia - Hypselodoris bennetti | | -2.865 | 0.004 | 0.325 | Aphelodoris varia - Hypselodoris bennetti | | -1.979 | 0.048 | 1 | Aphelodoris varia - Hypselodoris bennetti | | -3.351 | 0.001 | 0.063 |
| Chromodoris elisabethina - Hypselodoris bennetti | 1.54 | 0.124 | 1 | Chromodoris elisabethina - Hypselodoris bennetti | | 0.143 | 0.887 | 1 | Chromodoris elisabethina - Hypselodoris bennetti | | -1.403 | 0.161 | 1 | Chromodoris elisabethina - Hypselodoris bennetti | | -2.873 | 0.004 | 0.317 |
| Chromodoris kuiteri - Hypselodoris bennetti | 3.471 | 0.001 | 0.04 | Chromodoris kuiteri - Hypselodoris bennetti | | 3.021 | 0.003 | 0.196 | Chromodoris kuiteri - Hypselodoris bennetti | | 2.819 | 0.005 | 0.376 | Chromodoris kuiteri - Hypselodoris bennetti | | -3.074 | 0.002 | 0.165 |
| Discodoris sp - Hypselodoris bennetti | -1.144 | 0.253 | 1 | Discodoris sp - Hypselodoris bennetti | | -0.954 | 0.34 | 1 | Discodoris sp - Hypselodoris bennetti | | 0.504 | 0.614 | 1 | Discodoris sp - Hypselodoris bennetti | | -1.818 | 0.069 | 1 |
| Doriprismatica atromarginata - Hypselodoris bennetti | 2.281 | 0.023 | 1 | Doriprismatica atromarginata - Hypselodoris bennetti | | 0.08 | 0.936 | 1 | Doriprismatica atromarginata - Hypselodoris bennetti | | -1.02 | 0.308 | 1 | Doriprismatica atromarginata - Hypselodoris bennetti | | -2.598 | 0.009 | 0.732 |
| Glossodoris vespa - Hypselodoris bennetti | 0.869 | 0.385 | 1 | Glossodoris vespa - Hypselodoris bennetti | | 0.747 | 0.455 | 1 | Glossodoris vespa - Hypselodoris bennetti | | 1.141 | 0.254 | 1 | Glossodoris vespa - Hypselodoris bennetti | | -3.644 | <0.001 | 0.021 |
| Goniobranchus collingwoodi - Hypselodoris bennetti | -1.166 | 0.243 | 1 | Goniobranchus collingwoodi - Hypselodoris bennetti | | -1.808 | 0.071 | 1 | Goniobranchus collingwoodi - Hypselodoris bennetti | | -0.604 | 0.546 | 1 | Goniobranchus collingwoodi - Hypselodoris bennetti | | -2.997 | 0.003 | 0.213 |
| Goniobranchus splendidus - Hypselodoris bennetti | 2.835 | 0.005 | 0.357 | Goniobranchus splendidus - Hypselodoris bennetti | | 2.518 | 0.012 | 0.921 | Goniobranchus splendidus - Hypselodoris bennetti | | 2.096 | 0.036 | 1 | Goniobranchus splendidus - Hypselodoris bennetti | | -1.884 | 0.06 | 1 |
| Aphelodoris varia - Phyllidia elegans | 0.028 | 0.978 | 1 | Aphelodoris varia - Phyllidia elegans | | -0.158 | 0.875 | 1 | Aphelodoris varia - Phyllidia elegans | | -0.487 | 0.626 | 1 | Aphelodoris varia - Phyllidia elegans | | -1.364 | 0.172 | 1 |
| Chromodoris elisabethina - Phyllidia elegans | 3.673 | <0.001 | 0.019 | Chromodoris elisabethina - Phyllidia elegans | | 2.572 | 0.01 | 0.788 | Chromodoris elisabethina - Phyllidia elegans | | 0.017 | 0.987 | 1 | Chromodoris elisabethina - Phyllidia elegans | | -0.962 | 0.336 | 1 |
| Chromodoris kuiteri - Phyllidia elegans | 5.406 | <0.001 | <0.001 | Chromodoris kuiteri - Phyllidia elegans | | 5.173 | <0.001 | <0.001 | Chromodoris kuiteri - Phyllidia elegans | | 3.893 | <0.001 | 0.008 | Chromodoris kuiteri - Phyllidia elegans | | -1.214 | 0.225 | 1 |
| Discodoris sp - Phyllidia elegans | 1.068 | 0.286 | 1 | Discodoris sp - Phyllidia elegans | | 1.426 | 0.154 | 1 | Discodoris sp - Phyllidia elegans | | 1.718 | 0.086 | 1 | Discodoris sp - Phyllidia elegans | | -0.087 | 0.931 | 1 |
| Doriprismatica atromarginata - Phyllidia elegans | 4.419 | 0 | 0.001 | Doriprismatica atromarginata - Phyllidia elegans | | 2.593 | 0.01 | 0.743 | Doriprismatica atromarginata - Phyllidia elegans | | 0.419 | 0.675 | 1 | Doriprismatica atromarginata - Phyllidia elegans | | -0.639 | 0.523 | 1 |
| Glossodoris vespa - Phyllidia elegans | 2.945 | 0.003 | 0.252 | Glossodoris vespa - Phyllidia elegans | | 3.012 | 0.003 | 0.202 | Glossodoris vespa - Phyllidia elegans | | 2.311 | 0.021 | 1 | Glossodoris vespa - Phyllidia elegans | | -1.789 | 0.074 | 1 |
| Goniobranchus collingwoodi - Phyllidia elegans | 1.047 | 0.295 | 1 | Goniobranchus collingwoodi - Phyllidia elegans | | 0.63 | 0.529 | 1 | Goniobranchus collingwoodi - Phyllidia elegans | | 0.684 | 0.494 | 1 | Goniobranchus collingwoodi - Phyllidia elegans | | -1.186 | 0.236 | 1 |
| Goniobranchus splendidus - Phyllidia elegans | 4.912 | <0.001 | <0.001 | Goniobranchus splendidus - Phyllidia elegans | | 4.816 | <0.001 | <0.001 | Goniobranchus splendidus - Phyllidia elegans | | 3.275 | 0.001 | 0.082 | Goniobranchus splendidus - Phyllidia elegans | | -0.001 | 0.999 | 1 |
| Hypselodoris bennetti - Phyllidia elegans | 1.97 | 0.049 | 1 | Hypselodoris bennetti - Phyllidia elegans | | 2.138 | 0.033 | 1 | Hypselodoris bennetti - Phyllidia elegans | | 1.151 | 0.25 | 1 | Hypselodoris bennetti - Phyllidia elegans | | 1.485 | 0.138 | 1 |
| Aphelodoris varia - Phyllidia ocellata | -5 | <0.001 | <0.001 | Aphelodoris varia - Phyllidia ocellata | | -4.627 | <0.001 | <0.001 | Aphelodoris varia - Phyllidia ocellata | | -3.155 | 0.002 | 0.125 | Aphelodoris varia - Phyllidia ocellata | | -1.742 | 0.082 | 1 |
| Chromodoris elisabethina - Phyllidia ocellata | 0.184 | 0.854 | 1 | Chromodoris elisabethina - Phyllidia ocellata | | -0.719 | 0.472 | 1 | Chromodoris elisabethina - Phyllidia ocellata | | -2.369 | 0.018 | 1 | Chromodoris elisabethina - Phyllidia ocellata | | -1.163 | 0.245 | 1 |
| Chromodoris kuiteri - Phyllidia ocellata | 2.649 | 0.008 | 0.629 | Chromodoris kuiteri - Phyllidia ocellata | | 2.915 | 0.004 | 0.277 | Chromodoris kuiteri - Phyllidia ocellata | | 2.962 | 0.003 | 0.238 | Chromodoris kuiteri - Phyllidia ocellata | | -1.476 | 0.14 | 1 |
| Discodoris sp - Phyllidia ocellata | -3.022 | 0.003 | 0.196 | Discodoris sp - Phyllidia ocellata | | -1.992 | 0.046 | 1 | Discodoris sp - Phyllidia ocellata | | 0.091 | 0.928 | 1 | Discodoris sp - Phyllidia ocellata | | 0.032 | 0.974 | 1 |
| Doriprismatica atromarginata - Phyllidia ocellata | 1.087 | 0.277 | 1 | Doriprismatica atromarginata - Phyllidia ocellata | | -0.853 | 0.394 | 1 | Doriprismatica atromarginata - Phyllidia ocellata | | -1.95 | 0.051 | 1 | Doriprismatica atromarginata - Phyllidia ocellata | | -0.735 | 0.462 | 1 |
| Glossodoris vespa - Phyllidia ocellata | -0.546 | 0.585 | 1 | Glossodoris vespa - Phyllidia ocellata | | 0.1 | 0.92 | 1 | Glossodoris vespa - Phyllidia ocellata | | 0.874 | 0.382 | 1 | Glossodoris vespa - Phyllidia ocellata | | -2.213 | 0.027 | 1 |
| Goniobranchus collingwoodi - Phyllidia ocellata | -3.05 | 0.002 | 0.178 | Goniobranchus collingwoodi - Phyllidia ocellata | | -3.043 | 0.002 | 0.183 | Goniobranchus collingwoodi - Phyllidia ocellata | | -1.273 | 0.203 | 1 | Goniobranchus collingwoodi - Phyllidia ocellata | | -1.418 | 0.156 | 1 |
| Goniobranchus splendidus - Phyllidia ocellata | 1.816 | 0.069 | 1 | Goniobranchus splendidus - Phyllidia ocellata | | 2.32 | 0.02 | 1 | Goniobranchus splendidus - Phyllidia ocellata | | 2.106 | 0.035 | 1 | Goniobranchus splendidus - Phyllidia ocellata | | 0.167 | 0.867 | 1 |
| Hypselodoris bennetti - Phyllidia ocellata | -1.415 | 0.157 | 1 | Hypselodoris bennetti - Phyllidia ocellata | | -0.717 | 0.473 | 1 | Hypselodoris bennetti - Phyllidia ocellata | | -0.464 | 0.643 | 1 | Hypselodoris bennetti - Phyllidia ocellata | | 1.988 | 0.047 | 1 |
| Phyllidia elegans - Phyllidia ocellata | -3.583 | <0.001 | 0.027 | Phyllidia elegans - Phyllidia ocellata | | -3.132 | 0.002 | 0.135 | Phyllidia elegans - Phyllidia ocellata | | -1.759 | 0.079 | 1 | Phyllidia elegans - Phyllidia ocellata | | 0.119 | 0.906 | 1 |
| Aphelodoris varia - Phyllidia varicosa | -4.641 | 0 | 0 | Aphelodoris varia - Phyllidia varicosa | | -4.588 | <0.001 | <0.001 | Aphelodoris varia - Phyllidia varicosa | | -2.716 | 0.007 | 0.515 | Aphelodoris varia - Phyllidia varicosa | | 0.228 | 0.82 | 1 |
| Chromodoris elisabethina - Phyllidia varicosa | -0.914 | 0.361 | 1 | Chromodoris elisabethina - Phyllidia varicosa | | -1.781 | 0.075 | 1 | Chromodoris elisabethina - Phyllidia varicosa | | -2.174 | 0.03 | 1 | Chromodoris elisabethina - Phyllidia varicosa | | 0.603 | 0.547 | 1 |
| Chromodoris kuiteri - Phyllidia varicosa | 0.961 | 0.337 | 1 | Chromodoris kuiteri - Phyllidia varicosa | | 0.955 | 0.339 | 1 | Chromodoris kuiteri - Phyllidia varicosa | | 1.771 | 0.077 | 1 | Chromodoris kuiteri - Phyllidia varicosa | | 0.302 | 0.763 | 1 |
| Discodoris sp - Phyllidia varicosa | -3.286 | 0.001 | 0.079 | Discodoris sp - Phyllidia varicosa | | -2.704 | 0.007 | 0.534 | Discodoris sp - Phyllidia varicosa | | -0.361 | 0.718 | 1 | Discodoris sp - Phyllidia varicosa | | 1.398 | 0.162 | 1 |
| Doriprismatica atromarginata - Phyllidia varicosa | -0.316 | 0.752 | 1 | Doriprismatica atromarginata - Phyllidia varicosa | | -1.9 | 0.057 | 1 | Doriprismatica atromarginata - Phyllidia varicosa | | -1.842 | 0.065 | 1 | Doriprismatica atromarginata - Phyllidia varicosa | | 0.976 | 0.329 | 1 |
| Glossodoris vespa - Phyllidia varicosa | -1.409 | 0.159 | 1 | Glossodoris vespa - Phyllidia varicosa | | -1.118 | 0.263 | 1 | Glossodoris vespa - Phyllidia varicosa | | 0.233 | 0.816 | 1 | Glossodoris vespa - Phyllidia varicosa | | -0.305 | 0.761 | 1 |
| Goniobranchus collingwoodi - Phyllidia varicosa | -3.307 | 0.001 | 0.074 | Goniobranchus collingwoodi - Phyllidia varicosa | | -3.501 | <0.001 | 0.036 | Goniobranchus collingwoodi - Phyllidia varicosa | | -1.395 | 0.163 | 1 | Goniobranchus collingwoodi - Phyllidia varicosa | | 0.299 | 0.765 | 1 |
| Goniobranchus splendidus - Phyllidia varicosa | 0.22 | 0.826 | 1 | Goniobranchus splendidus - Phyllidia varicosa | | 0.363 | 0.716 | 1 | Goniobranchus splendidus - Phyllidia varicosa | | 1.035 | 0.301 | 1 | Goniobranchus splendidus - Phyllidia varicosa | | 1.599 | 0.11 | 1 |
| Hypselodoris bennetti - Phyllidia varicosa | -2.048 | 0.041 | 1 | Hypselodoris bennetti - Phyllidia varicosa | | -1.675 | 0.094 | 1 | Hypselodoris bennetti - Phyllidia varicosa | | -0.767 | 0.443 | 1 | Hypselodoris bennetti - Phyllidia varicosa | | 2.855 | 0.004 | 0.336 |
| Phyllidia elegans - Phyllidia varicosa | -3.812 | <0.001 | 0.011 | Phyllidia elegans - Phyllidia varicosa | | -3.617 | 0 | 0.023 | Phyllidia elegans - Phyllidia varicosa | | -1.82 | 0.069 | 1 | Phyllidia elegans - Phyllidia varicosa | | 1.3 | 0.194 | 1 |
| Phyllidia ocellata - Phyllidia varicosa | -1.061 | 0.289 | 1 | Phyllidia ocellata - Phyllidia varicosa | | -1.274 | 0.203 | 1 | Phyllidia ocellata - Phyllidia varicosa | | -0.458 | 0.647 | 1 | Phyllidia ocellata - Phyllidia varicosa | | 1.465 | 0.143 | 1 |
| Aphelodoris varia - Phyllidiella pustulosa | -1.446 | 0.148 | 1 | Aphelodoris varia - Phyllidiella pustulosa | | -1.092 | 0.275 | 1 | Aphelodoris varia - Phyllidiella pustulosa | | -0.482 | 0.63 | 1 | Aphelodoris varia - Phyllidiella pustulosa | | -1.927 | 0.054 | 1 |
| Chromodoris elisabethina - Phyllidiella pustulosa | 3.312 | 0.001 | 0.072 | Chromodoris elisabethina - Phyllidiella pustulosa | | 2.468 | 0.014 | 1 | Chromodoris elisabethina - Phyllidiella pustulosa | | 0.172 | 0.863 | 1 | Chromodoris elisabethina - Phyllidiella pustulosa | | -1.38 | 0.167 | 1 |
| Chromodoris kuiteri - Phyllidiella pustulosa | 5.487 | <0.001 | <0.001 | Chromodoris kuiteri - Phyllidiella pustulosa | | 5.742 | <0.001 | <0.001 | Chromodoris kuiteri - Phyllidiella pustulosa | | 5.079 | <0.001 | <0.001 | Chromodoris kuiteri - Phyllidiella pustulosa | | -1.669 | 0.095 | 1 |
| Discodoris sp - Phyllidiella pustulosa | 0.015 | 0.988 | 1 | Discodoris sp - Phyllidiella pustulosa | | 0.996 | 0.319 | 1 | Discodoris sp - Phyllidiella pustulosa | | 2.289 | 0.022 | 1 | Discodoris sp - Phyllidiella pustulosa | | -0.234 | 0.815 | 1 |
| Doriprismatica atromarginata - Phyllidiella pustulosa | 4.327 | <0.001 | 0.001 | Doriprismatica atromarginata - Phyllidiella pustulosa | | 2.522 | 0.012 | 0.909 | Doriprismatica atromarginata - Phyllidiella pustulosa | | 0.713 | 0.476 | 1 | Doriprismatica atromarginata - Phyllidiella pustulosa | | -0.989 | 0.322 | 1 |
| Glossodoris vespa - Phyllidiella pustulosa | 2.366 | 0.018 | 1 | Glossodoris vespa - Phyllidiella pustulosa | | 2.983 | 0.003 | 0.223 | Glossodoris vespa - Phyllidiella pustulosa | | 3.033 | 0.002 | 0.189 | Glossodoris vespa - Phyllidiella pustulosa | | -2.366 | 0.018 | 1 |
| Goniobranchus collingwoodi - Phyllidiella pustulosa | -0.011 | 0.991 | 1 | Goniobranchus collingwoodi - Phyllidiella pustulosa | | -0.001 | 0.999 | 1 | Goniobranchus collingwoodi - Phyllidiella pustulosa | | 0.995 | 0.32 | 1 | Goniobranchus collingwoodi - Phyllidiella pustulosa | | -1.61 | 0.107 | 1 |
| Goniobranchus splendidus - Phyllidiella pustulosa | 4.959 | <0.001 | <0.001 | Goniobranchus splendidus - Phyllidiella pustulosa | | 5.435 | <0.001 | <0.001 | Goniobranchus splendidus - Phyllidiella pustulosa | | 4.46 | <0.001 | 0.001 | Goniobranchus splendidus - Phyllidiella pustulosa | | -0.143 | 0.886 | 1 |
| Hypselodoris bennetti - Phyllidiella pustulosa | 1.197 | 0.231 | 1 | Hypselodoris bennetti - Phyllidiella pustulosa | | 1.871 | 0.061 | 1 | Hypselodoris bennetti - Phyllidiella pustulosa | | 1.507 | 0.132 | 1 | Hypselodoris bennetti - Phyllidiella pustulosa | | 1.675 | 0.094 | 1 |
| Phyllidia elegans - Phyllidiella pustulosa | -1.088 | 0.277 | 1 | Phyllidia elegans - Phyllidiella pustulosa | | -0.65 | 0.516 | 1 | Phyllidia elegans - Phyllidiella pustulosa | | 0.114 | 0.909 | 1 | Phyllidia elegans - Phyllidiella pustulosa | | -0.103 | 0.918 | 1 |
| Phyllidia ocellata - Phyllidiella pustulosa | 3.204 | 0.001 | 0.106 | Phyllidia ocellata - Phyllidiella pustulosa | | 3.208 | 0.001 | 0.104 | Phyllidia ocellata - Phyllidiella pustulosa | | 2.448 | 0.014 | 1 | Phyllidia ocellata - Phyllidiella pustulosa | | -0.294 | 0.769 | 1 |
| Phyllidia varicosa - Phyllidiella pustulosa | 3.398 | 0.001 | 0.053 | Phyllidia varicosa - Phyllidiella pustulosa | | 3.606 | <0.001 | 0.024 | Phyllidia varicosa - Phyllidiella pustulosa | | 2.256 | 0.024 | 1 | Phyllidia varicosa - Phyllidiella pustulosa | | -1.633 | 0.103 | 1 |

**Table S2.** Dunn tests at species level for chromatic detectability (Col.CoV). Significant (Bonferroni adjusted p) pairwise comparisons in red.

| **2cm** | | | | **5cm** | | | | **10cm** | | | | **30cm** | | | |
| --- | --- | --- | --- | --- | --- | --- | --- | --- | --- | --- | --- | --- | --- | --- | --- |
| Comparison | Z | P.unadj | P.adj | Comparison | Z | P.unadj | P.adj | Comparison | Z | P.unadj | P.adj | Comparison | Z | P.unadj | P.adj |
| Aphelodoris varia - Chromodoris elisabethina | -2.791 | 0.005 | 0.409 | Aphelodoris varia - Chromodoris elisabethina | -3.161 | 0.002 | 0.123 | Aphelodoris varia - Chromodoris elisabethina | -2.702 | 0.007 | 0.537 | Aphelodoris varia - Chromodoris elisabethina | 0.746 | 0.456 | 1 |
| Aphelodoris varia - Chromodoris kuiteri | -4.405 | <0.001 | 0.001 | Aphelodoris varia - Chromodoris kuiteri | -4.483 | <0.001 | 0.001 | Aphelodoris varia - Chromodoris kuiteri | -4.187 | <0.001 | 0.002 | Aphelodoris varia - Chromodoris kuiteri | -0.975 | 0.329 | 1 |
| Chromodoris elisabethina - Chromodoris kuiteri | -1.724 | 0.085 | 1 | Chromodoris elisabethina - Chromodoris kuiteri | -1.461 | 0.144 | 1 | Chromodoris elisabethina - Chromodoris kuiteri | -1.594 | 0.111 | 1 | Chromodoris elisabethina - Chromodoris kuiteri | -1.631 | 0.103 | 1 |
| Aphelodoris varia - Discodoris sp | 0.493 | 0.622 | 1 | Aphelodoris varia - Discodoris sp | 0.223 | 0.824 | 1 | Aphelodoris varia - Discodoris sp | 0.041 | 0.968 | 1 | Aphelodoris varia - Discodoris sp | 0.438 | 0.661 | 1 |
| Chromodoris elisabethina - Discodoris sp | 2.948 | 0.003 | 0.25 | Chromodoris elisabethina - Discodoris sp | 3.011 | 0.003 | 0.203 | Chromodoris elisabethina - Discodoris sp | 2.428 | 0.015 | 1 | Chromodoris elisabethina - Discodoris sp | -0.233 | 0.816 | 1 |
| Chromodoris kuiteri - Discodoris sp | 4.4 | <0.001 | 0.001 | Chromodoris kuiteri - Discodoris sp | 4.219 | <0.001 | 0.002 | Chromodoris kuiteri - Discodoris sp | 3.785 | <0.001 | 0.012 | Chromodoris kuiteri - Discodoris sp | 1.28 | 0.201 | 1 |
| Aphelodoris varia - Doriprismatica atromarginata | -1.724 | 0.085 | 1 | Aphelodoris varia - Doriprismatica atromarginata | -0.339 | 0.734 | 1 | Aphelodoris varia - Doriprismatica atromarginata | 0.474 | 0.636 | 1 | Aphelodoris varia - Doriprismatica atromarginata | 0.771 | 0.441 | 1 |
| Chromodoris elisabethina - Doriprismatica atromarginata | 1.205 | 0.228 | 1 | Chromodoris elisabethina - Doriprismatica atromarginata | 2.919 | 0.004 | 0.274 | Chromodoris elisabethina - Doriprismatica atromarginata | 3.232 | 0.001 | 0.096 | Chromodoris elisabethina - Doriprismatica atromarginata | -0.023 | 0.982 | 1 |
| Chromodoris kuiteri - Doriprismatica atromarginata | 2.948 | 0.003 | 0.249 | Chromodoris kuiteri - Doriprismatica atromarginata | 4.283 | <0.001 | 0.001 | Chromodoris kuiteri - Doriprismatica atromarginata | 4.717 | <0.001 | <0.001 | Chromodoris kuiteri - Doriprismatica atromarginata | 1.697 | 0.09 | 1 |
| Discodoris sp - Doriprismatica atromarginata | -2.006 | 0.045 | 1 | Discodoris sp - Doriprismatica atromarginata | -0.523 | 0.601 | 1 | Discodoris sp - Doriprismatica atromarginata | 0.371 | 0.711 | 1 | Discodoris sp - Doriprismatica atromarginata | 0.224 | 0.823 | 1 |
| Aphelodoris varia - Glossodoris vespa | -5.882 | <0.001 | <0.001 | Aphelodoris varia - Glossodoris vespa | -5.697 | <0.001 | <0.001 | Aphelodoris varia - Glossodoris vespa | -5.185 | <0.001 | <0.001 | Aphelodoris varia - Glossodoris vespa | -0.817 | 0.414 | 1 |
| Chromodoris elisabethina - Glossodoris vespa | -3.259 | 0.001 | 0.087 | Chromodoris elisabethina - Glossodoris vespa | -2.752 | 0.006 | 0.462 | Chromodoris elisabethina - Glossodoris vespa | -2.66 | 0.008 | 0.609 | Chromodoris elisabethina - Glossodoris vespa | -1.455 | 0.146 | 1 |
| Chromodoris kuiteri - Glossodoris vespa | -1.523 | 0.128 | 1 | Chromodoris kuiteri - Glossodoris vespa | -1.281 | 0.2 | 1 | Chromodoris kuiteri - Glossodoris vespa | -1.071 | 0.284 | 1 | Chromodoris kuiteri - Glossodoris vespa | 0.114 | 0.909 | 1 |
| Discodoris sp - Glossodoris vespa | -5.747 | 0 | 0 | Discodoris sp - Glossodoris vespa | -5.336 | 0 | 0 | Discodoris sp - Glossodoris vespa | -4.711 | 0 | 0 | Discodoris sp - Glossodoris vespa | -1.131 | 0.258 | 1 |
| Doriprismatica atromarginata - Glossodoris vespa | -4.51 | <0.001 | 0.001 | Doriprismatica atromarginata - Glossodoris vespa | -5.527 | <0.001 | <0.001 | Doriprismatica atromarginata - Glossodoris vespa | -5.712 | <0.001 | <0.001 | Doriprismatica atromarginata - Glossodoris vespa | -1.506 | 0.132 | 1 |
| Aphelodoris varia - Goniobranchus collingwoodi | -0.179 | 0.858 | 1 | Aphelodoris varia - Goniobranchus collingwoodi | -0.266 | 0.79 | 1 | Aphelodoris varia - Goniobranchus collingwoodi | -0.579 | 0.563 | 1 | Aphelodoris varia - Goniobranchus collingwoodi | 0.429 | 0.668 | 1 |
| Chromodoris elisabethina - Goniobranchus collingwoodi | 2.293 | 0.022 | 1 | Chromodoris elisabethina - Goniobranchus collingwoodi | 2.535 | 0.011 | 0.878 | Chromodoris elisabethina - Goniobranchus collingwoodi | 1.825 | 0.068 | 1 | Chromodoris elisabethina - Goniobranchus collingwoodi | -0.242 | 0.809 | 1 |
| Chromodoris kuiteri - Goniobranchus collingwoodi | 3.776 | <0.001 | 0.012 | Chromodoris kuiteri - Goniobranchus collingwoodi | 3.764 | <0.001 | 0.013 | Chromodoris kuiteri - Goniobranchus collingwoodi | 3.209 | 0.001 | 0.104 | Chromodoris kuiteri - Goniobranchus collingwoodi | 1.271 | 0.204 | 1 |
| Discodoris sp - Goniobranchus collingwoodi | -0.606 | 0.545 | 1 | Discodoris sp - Goniobranchus collingwoodi | -0.441 | 0.659 | 1 | Discodoris sp - Goniobranchus collingwoodi | -0.558 | 0.577 | 1 | Discodoris sp - Goniobranchus collingwoodi | -0.008 | 0.993 | 1 |
| Doriprismatica atromarginata - Goniobranchus collingwoodi | 1.319 | 0.187 | 1 | Doriprismatica atromarginata - Goniobranchus collingwoodi | 0.023 | 0.981 | 1 | Doriprismatica atromarginata - Goniobranchus collingwoodi | -1.004 | 0.315 | 1 | Doriprismatica atromarginata - Goniobranchus collingwoodi | -0.234 | 0.815 | 1 |
| Glossodoris vespa - Goniobranchus collingwoodi | 5.141 | 0 | 0 | Glossodoris vespa - Goniobranchus collingwoodi | 4.895 | 0 | 0 | Glossodoris vespa - Goniobranchus collingwoodi | 4.152 | 0 | 0.003 | Glossodoris vespa - Goniobranchus collingwoodi | 1.123 | 0.262 | 1 |
| Aphelodoris varia - Goniobranchus splendidus | -3.82 | <0.001 | 0.01 | Aphelodoris varia - Goniobranchus splendidus | -3.636 | <0.001 | 0.022 | Aphelodoris varia - Goniobranchus splendidus | -2.136 | 0.033 | 1 | Aphelodoris varia - Goniobranchus splendidus | 0.409 | 0.683 | 1 |
| Chromodoris elisabethina - Goniobranchus splendidus | -0.87 | 0.384 | 1 | Chromodoris elisabethina - Goniobranchus splendidus | -0.319 | 0.75 | 1 | Chromodoris elisabethina - Goniobranchus splendidus | 0.665 | 0.506 | 1 | Chromodoris elisabethina - Goniobranchus splendidus | -0.359 | 0.72 | 1 |
| Chromodoris kuiteri - Goniobranchus splendidus | 0.97 | 0.332 | 1 | Chromodoris kuiteri - Goniobranchus splendidus | 1.216 | 0.224 | 1 | Chromodoris kuiteri - Goniobranchus splendidus | 2.281 | 0.023 | 1 | Chromodoris kuiteri - Goniobranchus splendidus | 1.355 | 0.175 | 1 |
| Discodoris sp - Goniobranchus splendidus | -3.839 | <0.001 | 0.01 | Discodoris sp - Goniobranchus splendidus | -3.406 | 0.001 | 0.051 | Discodoris sp - Goniobranchus splendidus | -1.91 | 0.056 | 1 | Discodoris sp - Goniobranchus splendidus | -0.084 | 0.933 | 1 |
| Doriprismatica atromarginata - Goniobranchus splendidus | -2.19 | 0.029 | 1 | Doriprismatica atromarginata - Goniobranchus splendidus | -3.4 | 0.001 | 0.052 | Doriprismatica atromarginata - Goniobranchus splendidus | -2.678 | 0.007 | 0.578 | Doriprismatica atromarginata - Goniobranchus splendidus | -0.358 | 0.72 | 1 |
| Glossodoris vespa - Goniobranchus splendidus | 2.586 | 0.01 | 0.758 | Glossodoris vespa - Goniobranchus splendidus | 2.56 | 0.01 | 0.817 | Glossodoris vespa - Goniobranchus splendidus | 3.357 | 0.001 | 0.062 | Glossodoris vespa - Goniobranchus splendidus | 1.181 | 0.238 | 1 |
| Goniobranchus collingwoodi - Goniobranchus splendidus | -3.162 | 0.002 | 0.122 | Goniobranchus collingwoodi - Goniobranchus splendidus | -2.913 | 0.004 | 0.279 | Goniobranchus collingwoodi - Goniobranchus splendidus | -1.286 | 0.199 | 1 | Goniobranchus collingwoodi - Goniobranchus splendidus | -0.074 | 0.941 | 1 |
| Aphelodoris varia - Hypselodoris bennetti | -4.041 | <0.001 | 0.004 | Aphelodoris varia - Hypselodoris bennetti | -3.626 | <0.001 | 0.022 | Aphelodoris varia - Hypselodoris bennetti | -1.962 | 0.05 | 1 | Aphelodoris varia - Hypselodoris bennetti | -1.022 | 0.307 | 1 |
| Chromodoris elisabethina - Hypselodoris bennetti | -1.788 | 0.074 | 1 | Chromodoris elisabethina - Hypselodoris bennetti | -1.094 | 0.274 | 1 | Chromodoris elisabethina - Hypselodoris bennetti | 0.179 | 0.858 | 1 | Chromodoris elisabethina - Hypselodoris bennetti | -1.581 | 0.114 | 1 |
| Chromodoris kuiteri - Hypselodoris bennetti | -0.312 | 0.755 | 1 | Chromodoris kuiteri - Hypselodoris bennetti | 0.142 | 0.887 | 1 | Chromodoris kuiteri - Hypselodoris bennetti | 1.477 | 0.14 | 1 | Chromodoris kuiteri - Hypselodoris bennetti | -0.189 | 0.85 | 1 |
| Discodoris sp - Hypselodoris bennetti | -4.123 | <0.001 | 0.003 | Discodoris sp - Hypselodoris bennetti | -3.523 | <0.001 | 0.033 | Discodoris sp - Hypselodoris bennetti | -1.842 | 0.065 | 1 | Discodoris sp - Hypselodoris bennetti | -1.295 | 0.195 | 1 |
| Doriprismatica atromarginata - Hypselodoris bennetti | -2.802 | 0.005 | 0.396 | Doriprismatica atromarginata - Hypselodoris bennetti | -3.43 | 0.001 | 0.047 | Doriprismatica atromarginata - Hypselodoris bennetti | -2.354 | 0.019 | 1 | Doriprismatica atromarginata - Hypselodoris bennetti | -1.623 | 0.105 | 1 |
| Glossodoris vespa - Hypselodoris bennetti | 1.016 | 0.309 | 1 | Glossodoris vespa - Hypselodoris bennetti | 1.25 | 0.211 | 1 | Glossodoris vespa - Hypselodoris bennetti | 2.371 | 0.018 | 1 | Glossodoris vespa - Hypselodoris bennetti | -0.283 | 0.777 | 1 |
| Goniobranchus collingwoodi - Hypselodoris bennetti | -3.581 | <0.001 | 0.027 | Goniobranchus collingwoodi - Hypselodoris bennetti | -3.128 | 0.002 | 0.137 | Goniobranchus collingwoodi - Hypselodoris bennetti | -1.342 | 0.179 | 1 | Goniobranchus collingwoodi - Hypselodoris bennetti | -1.287 | 0.198 | 1 |
| Goniobranchus splendidus - Hypselodoris bennetti | -1.148 | 0.251 | 1 | Goniobranchus splendidus - Hypselodoris bennetti | -0.871 | 0.384 | 1 | Goniobranchus splendidus - Hypselodoris bennetti | -0.343 | 0.732 | 1 | Goniobranchus splendidus - Hypselodoris bennetti | -1.34 | 0.18 | 1 |
| Aphelodoris varia - Phyllidia elegans | 0.562 | 0.574 | 1 | Aphelodoris varia - Phyllidia elegans | 0.826 | 0.409 | 1 | Aphelodoris varia - Phyllidia elegans | 0.982 | 0.326 | 1 | Aphelodoris varia - Phyllidia elegans | 1.319 | 0.187 | 1 |
| Chromodoris elisabethina - Phyllidia elegans | 2.56 | 0.01 | 0.817 | Chromodoris elisabethina - Phyllidia elegans | 3.085 | 0.002 | 0.159 | Chromodoris elisabethina - Phyllidia elegans | 2.908 | 0.004 | 0.284 | Chromodoris elisabethina - Phyllidia elegans | 0.759 | 0.448 | 1 |
| Chromodoris kuiteri - Phyllidia elegans | 3.792 | <0.001 | 0.012 | Chromodoris kuiteri - Phyllidia elegans | 4.101 | <0.001 | 0.003 | Chromodoris kuiteri - Phyllidia elegans | 4.031 | <0.001 | 0.004 | Chromodoris kuiteri - Phyllidia elegans | 1.977 | 0.048 | 1 |
| Discodoris sp - Phyllidia elegans | 0.153 | 0.878 | 1 | Discodoris sp - Phyllidia elegans | 0.602 | 0.547 | 1 | Discodoris sp - Phyllidia elegans | 0.885 | 0.376 | 1 | Discodoris sp - Phyllidia elegans | 0.901 | 0.368 | 1 |
| Doriprismatica atromarginata - Phyllidia elegans | 1.771 | 0.077 | 1 | Doriprismatica atromarginata - Phyllidia elegans | 1.074 | 0.283 | 1 | Doriprismatica atromarginata - Phyllidia elegans | 0.666 | 0.506 | 1 | Doriprismatica atromarginata - Phyllidia elegans | 0.8 | 0.424 | 1 |
| Glossodoris vespa - Phyllidia elegans | 4.946 | 0 | 0 | Glossodoris vespa - Phyllidia elegans | 5.053 | 0 | 0 | Glossodoris vespa - Phyllidia elegans | 4.814 | 0 | 0 | Glossodoris vespa - Phyllidia elegans | 1.844 | 0.065 | 1 |
| Goniobranchus collingwoodi - Phyllidia elegans | 0.659 | 0.51 | 1 | Goniobranchus collingwoodi - Phyllidia elegans | 0.97 | 0.332 | 1 | Goniobranchus collingwoodi - Phyllidia elegans | 1.351 | 0.177 | 1 | Goniobranchus collingwoodi - Phyllidia elegans | 0.908 | 0.364 | 1 |
| Goniobranchus splendidus - Phyllidia elegans | 3.252 | 0.001 | 0.089 | Goniobranchus splendidus - Phyllidia elegans | 3.388 | 0.001 | 0.055 | Goniobranchus splendidus - Phyllidia elegans | 2.49 | 0.013 | 0.998 | Goniobranchus splendidus - Phyllidia elegans | 1.038 | 0.299 | 1 |
| Hypselodoris bennetti - Phyllidia elegans | 3.69 | <0.001 | 0.017 | Hypselodoris bennetti - Phyllidia elegans | 3.588 | <0.001 | 0.026 | Hypselodoris bennetti - Phyllidia elegans | 2.402 | 0.016 | 1 | Hypselodoris bennetti - Phyllidia elegans | 1.946 | 0.052 | 1 |
| Aphelodoris varia - Phyllidia ocellata | 0.62 | 0.535 | 1 | Aphelodoris varia - Phyllidia ocellata | 0.792 | 0.429 | 1 | Aphelodoris varia - Phyllidia ocellata | 0.902 | 0.367 | 1 | Aphelodoris varia - Phyllidia ocellata | 2.16 | 0.031 | 1 |
| Chromodoris elisabethina - Phyllidia ocellata | 3.363 | 0.001 | 0.06 | Chromodoris elisabethina - Phyllidia ocellata | 3.895 | 0 | 0.008 | Chromodoris elisabethina - Phyllidia ocellata | 3.547 | 0 | 0.03 | Chromodoris elisabethina - Phyllidia ocellata | 1.35 | 0.177 | 1 |
| Chromodoris kuiteri - Phyllidia ocellata | 4.932 | <0.001 | <0.001 | Chromodoris kuiteri - Phyllidia ocellata | 5.165 | <0.001 | <0.001 | Chromodoris kuiteri - Phyllidia ocellata | 4.973 | <0.001 | <0.001 | Chromodoris kuiteri - Phyllidia ocellata | 2.938 | 0.003 | 0.258 |
| Discodoris sp - Phyllidia ocellata | 0.056 | 0.955 | 1 | Discodoris sp - Phyllidia ocellata | 0.475 | 0.635 | 1 | Discodoris sp - Phyllidia ocellata | 0.753 | 0.452 | 1 | Discodoris sp - Phyllidia ocellata | 1.465 | 0.143 | 1 |
| Doriprismatica atromarginata - Phyllidia ocellata | 2.342 | 0.019 | 1 | Doriprismatica atromarginata - Phyllidia ocellata | 1.149 | 0.25 | 1 | Doriprismatica atromarginata - Phyllidia ocellata | 0.459 | 0.646 | 1 | Doriprismatica atromarginata - Phyllidia ocellata | 1.459 | 0.144 | 1 |
| Glossodoris vespa - Phyllidia ocellata | 6.379 | <0.001 | <0.001 | Glossodoris vespa - Phyllidia ocellata | 6.346 | <0.001 | <0.001 | Glossodoris vespa - Phyllidia ocellata | 5.935 | <0.001 | <0.001 | Glossodoris vespa - Phyllidia ocellata | 2.709 | 0.007 | 0.526 |
| Goniobranchus collingwoodi - Phyllidia ocellata | 0.723 | 0.47 | 1 | Goniobranchus collingwoodi - Phyllidia ocellata | 0.96 | 0.337 | 1 | Goniobranchus collingwoodi - Phyllidia ocellata | 1.367 | 0.172 | 1 | Goniobranchus collingwoodi - Phyllidia ocellata | 1.474 | 0.14 | 1 |
| Goniobranchus splendidus - Phyllidia ocellata | 4.404 | <0.001 | 0.001 | Goniobranchus splendidus - Phyllidia ocellata | 4.395 | <0.001 | 0.001 | Goniobranchus splendidus - Phyllidia ocellata | 3.024 | 0.002 | 0.195 | Goniobranchus splendidus - Phyllidia ocellata | 1.778 | 0.075 | 1 |
| Hypselodoris bennetti - Phyllidia ocellata | 4.493 | <0.001 | 0.001 | Hypselodoris bennetti - Phyllidia ocellata | 4.213 | <0.001 | 0.002 | Hypselodoris bennetti - Phyllidia ocellata | 2.645 | 0.008 | 0.638 | Hypselodoris bennetti - Phyllidia ocellata | 2.679 | 0.007 | 0.575 |
| Phyllidia elegans - Phyllidia ocellata | -0.118 | 0.906 | 1 | Phyllidia elegans - Phyllidia ocellata | -0.259 | 0.796 | 1 | Phyllidia elegans - Phyllidia ocellata | -0.335 | 0.737 | 1 | Phyllidia elegans - Phyllidia ocellata | 0.224 | 0.823 | 1 |
| Aphelodoris varia - Phyllidia varicosa | -1.236 | 0.216 | 1 | Aphelodoris varia - Phyllidia varicosa | -1.773 | 0.076 | 1 | Aphelodoris varia - Phyllidia varicosa | -1.608 | 0.108 | 1 | Aphelodoris varia - Phyllidia varicosa | 0.701 | 0.483 | 1 |
| Chromodoris elisabethina - Phyllidia varicosa | 0.793 | 0.428 | 1 | Chromodoris elisabethina - Phyllidia varicosa | 0.531 | 0.595 | 1 | Chromodoris elisabethina - Phyllidia varicosa | 0.364 | 0.716 | 1 | Chromodoris elisabethina - Phyllidia varicosa | 0.152 | 0.879 | 1 |
| Chromodoris kuiteri - Phyllidia varicosa | 2.08 | 0.038 | 1 | Chromodoris kuiteri - Phyllidia varicosa | 1.626 | 0.104 | 1 | Chromodoris kuiteri - Phyllidia varicosa | 1.565 | 0.118 | 1 | Chromodoris kuiteri - Phyllidia varicosa | 1.389 | 0.165 | 1 |
| Discodoris sp - Phyllidia varicosa | -1.524 | 0.128 | 1 | Discodoris sp - Phyllidia varicosa | -1.821 | 0.069 | 1 | Discodoris sp - Phyllidia varicosa | -1.53 | 0.126 | 1 | Discodoris sp - Phyllidia varicosa | 0.324 | 0.746 | 1 |
| Doriprismatica atromarginata - Phyllidia varicosa | -0.052 | 0.958 | 1 | Doriprismatica atromarginata - Phyllidia varicosa | -1.562 | 0.118 | 1 | Doriprismatica atromarginata - Phyllidia varicosa | -1.961 | 0.05 | 1 | Doriprismatica atromarginata - Phyllidia varicosa | 0.173 | 0.862 | 1 |
| Glossodoris vespa - Phyllidia varicosa | 3.269 | 0.001 | 0.084 | Glossodoris vespa - Phyllidia varicosa | 2.629 | 0.009 | 0.667 | Glossodoris vespa - Phyllidia varicosa | 2.399 | 0.016 | 1 | Glossodoris vespa - Phyllidia varicosa | 1.268 | 0.205 | 1 |
| Goniobranchus collingwoodi - Phyllidia varicosa | -1.018 | 0.309 | 1 | Goniobranchus collingwoodi - Phyllidia varicosa | -1.453 | 0.146 | 1 | Goniobranchus collingwoodi - Phyllidia varicosa | -1.064 | 0.287 | 1 | Goniobranchus collingwoodi - Phyllidia varicosa | 0.331 | 0.74 | 1 |
| Goniobranchus splendidus - Phyllidia varicosa | 1.445 | 0.149 | 1 | Goniobranchus splendidus - Phyllidia varicosa | 0.776 | 0.438 | 1 | Goniobranchus splendidus - Phyllidia varicosa | -0.113 | 0.91 | 1 | Goniobranchus splendidus - Phyllidia varicosa | 0.417 | 0.677 | 1 |
| Hypselodoris bennetti - Phyllidia varicosa | 2.143 | 0.032 | 1 | Hypselodoris bennetti - Phyllidia varicosa | 1.351 | 0.177 | 1 | Hypselodoris bennetti - Phyllidia varicosa | 0.173 | 0.862 | 1 | Hypselodoris bennetti - Phyllidia varicosa | 1.414 | 0.157 | 1 |
| Phyllidia elegans - Phyllidia varicosa | -1.468 | 0.142 | 1 | Phyllidia elegans - Phyllidia varicosa | -2.122 | 0.034 | 1 | Phyllidia elegans - Phyllidia varicosa | -2.114 | 0.034 | 1 | Phyllidia elegans - Phyllidia varicosa | -0.505 | 0.614 | 1 |
| Phyllidia ocellata - Phyllidia varicosa | -1.671 | 0.095 | 1 | Phyllidia ocellata - Phyllidia varicosa | -2.326 | 0.02 | 1 | Phyllidia ocellata - Phyllidia varicosa | -2.24 | 0.025 | 1 | Phyllidia ocellata - Phyllidia varicosa | -0.839 | 0.402 | 1 |
| Aphelodoris varia - Phyllidiella pustulosa | -1.451 | 0.147 | 1 | Aphelodoris varia - Phyllidiella pustulosa | -1.442 | 0.149 | 1 | Aphelodoris varia - Phyllidiella pustulosa | -0.818 | 0.413 | 1 | Aphelodoris varia - Phyllidiella pustulosa | 0.017 | 0.986 | 1 |
| Chromodoris elisabethina - Phyllidiella pustulosa | 1.188 | 0.235 | 1 | Chromodoris elisabethina - Phyllidiella pustulosa | 1.541 | 0.123 | 1 | Chromodoris elisabethina - Phyllidiella pustulosa | 1.719 | 0.086 | 1 | Chromodoris elisabethina - Phyllidiella pustulosa | -0.678 | 0.498 | 1 |
| Chromodoris kuiteri - Phyllidiella pustulosa | 2.791 | 0.005 | 0.409 | Chromodoris kuiteri - Phyllidiella pustulosa | 2.873 | 0.004 | 0.317 | Chromodoris kuiteri - Phyllidiella pustulosa | 3.171 | 0.002 | 0.119 | Chromodoris kuiteri - Phyllidiella pustulosa | 0.93 | 0.352 | 1 |
| Discodoris sp - Phyllidiella pustulosa | -1.759 | 0.079 | 1 | Discodoris sp - Phyllidiella pustulosa | -1.496 | 0.135 | 1 | Discodoris sp - Phyllidiella pustulosa | -0.768 | 0.442 | 1 | Discodoris sp - Phyllidiella pustulosa | -0.397 | 0.691 | 1 |
| Doriprismatica atromarginata - Phyllidiella pustulosa | 0.102 | 0.918 | 1 | Doriprismatica atromarginata - Phyllidiella pustulosa | -1.164 | 0.244 | 1 | Doriprismatica atromarginata - Phyllidiella pustulosa | -1.275 | 0.202 | 1 | Doriprismatica atromarginata - Phyllidiella pustulosa | -0.693 | 0.488 | 1 |
| Glossodoris vespa - Phyllidiella pustulosa | 4.243 | <0.001 | 0.002 | Glossodoris vespa - Phyllidiella pustulosa | 4.078 | <0.001 | 0.004 | Glossodoris vespa - Phyllidiella pustulosa | 4.152 | <0.001 | 0.003 | Glossodoris vespa - Phyllidiella pustulosa | 0.784 | 0.433 | 1 |
| Goniobranchus collingwoodi - Phyllidiella pustulosa | -1.126 | 0.26 | 1 | Goniobranchus collingwoodi - Phyllidiella pustulosa | -1.035 | 0.301 | 1 | Goniobranchus collingwoodi - Phyllidiella pustulosa | -0.185 | 0.853 | 1 | Goniobranchus collingwoodi - Phyllidiella pustulosa | -0.388 | 0.698 | 1 |
| Goniobranchus splendidus - Phyllidiella pustulosa | 2.067 | 0.039 | 1 | Goniobranchus splendidus - Phyllidiella pustulosa | 1.907 | 0.057 | 1 | Goniobranchus splendidus - Phyllidiella pustulosa | 1.149 | 0.25 | 1 | Goniobranchus splendidus - Phyllidiella pustulosa | -0.361 | 0.718 | 1 |
| Hypselodoris bennetti - Phyllidiella pustulosa | 2.709 | 0.007 | 0.526 | Hypselodoris bennetti - Phyllidiella pustulosa | 2.321 | 0.02 | 1 | Hypselodoris bennetti - Phyllidiella pustulosa | 1.226 | 0.22 | 1 | Hypselodoris bennetti - Phyllidiella pustulosa | 0.988 | 0.323 | 1 |
| Phyllidia elegans - Phyllidiella pustulosa | -1.605 | 0.109 | 1 | Phyllidia elegans - Phyllidiella pustulosa | -1.851 | 0.064 | 1 | Phyllidia elegans - Phyllidiella pustulosa | -1.544 | 0.123 | 1 | Phyllidia elegans - Phyllidiella pustulosa | -1.255 | 0.21 | 1 |
| Phyllidia ocellata - Phyllidiella pustulosa | -2.013 | 0.044 | 1 | Phyllidia ocellata - Phyllidiella pustulosa | -2.162 | 0.031 | 1 | Phyllidia ocellata - Phyllidiella pustulosa | -1.647 | 0.1 | 1 | Phyllidia ocellata - Phyllidiella pustulosa | -1.986 | 0.047 | 1 |
| Phyllidia varicosa - Phyllidiella pustulosa | 0.123 | 0.902 | 1 | Phyllidia varicosa - Phyllidiella pustulosa | 0.646 | 0.518 | 1 | Phyllidia varicosa - Phyllidiella pustulosa | 0.944 | 0.345 | 1 | Phyllidia varicosa - Phyllidiella pustulosa | -0.661 | 0.509 | 1 |

**Table S3.** Dunn tests at species level for achromatic boldness (Lum.CoV). Significant (Bonferroni adjusted p) pairwise comparisons in red.

| **2cm** | | | | **5cm** | | | | **10cm** | | | | **30cm** | | | |
| --- | --- | --- | --- | --- | --- | --- | --- | --- | --- | --- | --- | --- | --- | --- | --- |
| Comparison | Z | P.unadj | P.adj | Comparison | Z | P.unadj | P.adj | Comparison | Z | P.unadj | P.adj | Comparison | Z | P.unadj | P.adj |
| Aphelodoris varia - Chromodoris elisabethina | -3.159 | 0.002 | 0.123 | Aphelodoris varia - Chromodoris elisabethina | -2.532 | 0.011 | 0.884 | Aphelodoris varia - Chromodoris elisabethina | -1.323 | 0.186 | 1 | Aphelodoris varia - Chromodoris elisabethina | 0.315 | 0.753 | 1 |
| Aphelodoris varia - Chromodoris kuiteri | -5.621 | <0.001 | <0.001 | Aphelodoris varia - Chromodoris kuiteri | -5.761 | <0.001 | <0.001 | Aphelodoris varia - Chromodoris kuiteri | -5.28 | <0.001 | <0.001 | Aphelodoris varia - Chromodoris kuiteri | -2.921 | 0.003 | 0.272 |
| Chromodoris elisabethina - Chromodoris kuiteri | -2.568 | 0.01 | 0.797 | Chromodoris elisabethina - Chromodoris kuiteri | -3.279 | 0.001 | 0.081 | Chromodoris elisabethina - Chromodoris kuiteri | -3.919 | 0 | 0.007 | Chromodoris elisabethina - Chromodoris kuiteri | -3.127 | 0.002 | 0.138 |
| Aphelodoris varia - Discodoris sp | 1.799 | 0.072 | 1 | Aphelodoris varia - Discodoris sp | 2.027 | 0.043 | 1 | Aphelodoris varia - Discodoris sp | 2.093 | 0.036 | 1 | Aphelodoris varia - Discodoris sp | 0.3 | 0.764 | 1 |
| Chromodoris elisabethina - Discodoris sp | 4.543 | <0.001 | <0.001 | Chromodoris elisabethina - Discodoris sp | 4.212 | <0.001 | 0.002 | Chromodoris elisabethina - Discodoris sp | 3.207 | 0.001 | 0.104 | Chromodoris elisabethina - Discodoris sp | 0.013 | 0.989 | 1 |
| Chromodoris kuiteri - Discodoris sp | 6.701 | <0.001 | <0.001 | Chromodoris kuiteri - Discodoris sp | 7.039 | <0.001 | <0.001 | Chromodoris kuiteri - Discodoris sp | 6.67 | <0.001 | <0.001 | Chromodoris kuiteri - Discodoris sp | 2.893 | 0.004 | 0.298 |
| Aphelodoris varia - Doriprismatica atromarginata | -4.149 | <0.001 | 0.003 | Aphelodoris varia - Doriprismatica atromarginata | -2.336 | 0.019 | 1 | Aphelodoris varia - Doriprismatica atromarginata | 0.028 | 0.978 | 1 | Aphelodoris varia - Doriprismatica atromarginata | 1.794 | 0.073 | 1 |
| Chromodoris elisabethina - Doriprismatica atromarginata | -0.756 | 0.45 | 1 | Chromodoris elisabethina - Doriprismatica atromarginata | 0.348 | 0.728 | 1 | Chromodoris elisabethina - Doriprismatica atromarginata | 1.385 | 0.166 | 1 | Chromodoris elisabethina - Doriprismatica atromarginata | 1.407 | 0.16 | 1 |
| Chromodoris kuiteri - Doriprismatica atromarginata | 1.996 | 0.046 | 1 | Chromodoris kuiteri - Doriprismatica atromarginata | 3.782 | <0.001 | 0.012 | Chromodoris kuiteri - Doriprismatica atromarginata | 5.431 | <0.001 | <0.001 | Chromodoris kuiteri - Doriprismatica atromarginata | 4.617 | 0 | 0 |
| Discodoris sp - Doriprismatica atromarginata | -5.453 | <0.001 | <0.001 | Discodoris sp - Doriprismatica atromarginata | -4.107 | <0.001 | 0.003 | Discodoris sp - Doriprismatica atromarginata | -2.115 | 0.034 | 1 | Discodoris sp - Doriprismatica atromarginata | 1.257 | 0.209 | 1 |
| Aphelodoris varia - Glossodoris vespa | -2.712 | 0.007 | 0.522 | Aphelodoris varia - Glossodoris vespa | -3.453 | 0.001 | 0.043 | Aphelodoris varia - Glossodoris vespa | -3.198 | 0.001 | 0.108 | Aphelodoris varia - Glossodoris vespa | -0.143 | 0.886 | 1 |
| Chromodoris elisabethina - Glossodoris vespa | 0.152 | 0.879 | 1 | Chromodoris elisabethina - Glossodoris vespa | -1.124 | 0.261 | 1 | Chromodoris elisabethina - Glossodoris vespa | -1.944 | 0.052 | 1 | Chromodoris elisabethina - Glossodoris vespa | -0.418 | 0.676 | 1 |
| Chromodoris kuiteri - Glossodoris vespa | 2.511 | 0.012 | 0.94 | Chromodoris kuiteri - Glossodoris vespa | 1.947 | 0.051 | 1 | Chromodoris kuiteri - Glossodoris vespa | 1.754 | 0.079 | 1 | Chromodoris kuiteri - Glossodoris vespa | 2.481 | 0.013 | 1 |
| Discodoris sp - Glossodoris vespa | -4.066 | <0.001 | 0.004 | Discodoris sp - Glossodoris vespa | -4.94 | <0.001 | <0.001 | Discodoris sp - Glossodoris vespa | -4.769 | <0.001 | <0.001 | Discodoris sp - Glossodoris vespa | -0.399 | 0.69 | 1 |
| Doriprismatica atromarginata - Glossodoris vespa | 0.843 | 0.399 | 1 | Doriprismatica atromarginata - Glossodoris vespa | -1.494 | 0.135 | 1 | Doriprismatica atromarginata - Glossodoris vespa | -3.292 | 0.001 | 0.077 | Doriprismatica atromarginata - Glossodoris vespa | -1.71 | 0.087 | 1 |
| Aphelodoris varia - Goniobranchus collingwoodi | 2.387 | 0.017 | 1 | Aphelodoris varia - Goniobranchus collingwoodi | 2.188 | 0.029 | 1 | Aphelodoris varia - Goniobranchus collingwoodi | 1.997 | 0.046 | 1 | Aphelodoris varia - Goniobranchus collingwoodi | -0.326 | 0.744 | 1 |
| Chromodoris elisabethina - Goniobranchus collingwoodi | 5.116 | <0.001 | <0.001 | Chromodoris elisabethina - Goniobranchus collingwoodi | 4.369 | <0.001 | 0.001 | Chromodoris elisabethina - Goniobranchus collingwoodi | 3.114 | 0.002 | 0.144 | Chromodoris elisabethina - Goniobranchus collingwoodi | -0.596 | 0.551 | 1 |
| Chromodoris kuiteri - Goniobranchus collingwoodi | 7.248 | <0.001 | <0.001 | Chromodoris kuiteri - Goniobranchus collingwoodi | 7.189 | <0.001 | <0.001 | Chromodoris kuiteri - Goniobranchus collingwoodi | 6.581 | <0.001 | <0.001 | Chromodoris kuiteri - Goniobranchus collingwoodi | 2.311 | 0.021 | 1 |
| Discodoris sp - Goniobranchus collingwoodi | 0.531 | 0.596 | 1 | Discodoris sp - Goniobranchus collingwoodi | 0.145 | 0.885 | 1 | Discodoris sp - Goniobranchus collingwoodi | -0.087 | 0.931 | 1 | Discodoris sp - Goniobranchus collingwoodi | -0.564 | 0.573 | 1 |
| Doriprismatica atromarginata - Goniobranchus collingwoodi | 6.054 | 0 | 0 | Doriprismatica atromarginata - Goniobranchus collingwoodi | 4.272 | <0.001 | 0.002 | Doriprismatica atromarginata - Goniobranchus collingwoodi | 2.017 | 0.044 | 1 | Doriprismatica atromarginata - Goniobranchus collingwoodi | -1.897 | 0.058 | 1 |
| Glossodoris vespa - Goniobranchus collingwoodi | 4.596 | 0 | 0 | Glossodoris vespa - Goniobranchus collingwoodi | 5.085 | <0.001 | <0.001 | Glossodoris vespa - Goniobranchus collingwoodi | 4.683 | <0.001 | <0.001 | Glossodoris vespa - Goniobranchus collingwoodi | -0.165 | 0.869 | 1 |
| Aphelodoris varia - Goniobranchus splendidus | -4.401 | 0 | 0.001 | Aphelodoris varia - Goniobranchus splendidus | -4.435 | <0.001 | 0.001 | Aphelodoris varia - Goniobranchus splendidus | -3.582 | <0.001 | 0.027 | Aphelodoris varia - Goniobranchus splendidus | 0.229 | 0.819 | 1 |
| Chromodoris elisabethina - Goniobranchus splendidus | -1.059 | 0.289 | 1 | Chromodoris elisabethina - Goniobranchus splendidus | -1.726 | 0.084 | 1 | Chromodoris elisabethina - Goniobranchus splendidus | -2.122 | 0.034 | 1 | Chromodoris elisabethina - Goniobranchus splendidus | -0.097 | 0.923 | 1 |
| Chromodoris kuiteri - Goniobranchus splendidus | 1.668 | 0.095 | 1 | Chromodoris kuiteri - Goniobranchus splendidus | 1.778 | 0.075 | 1 | Chromodoris kuiteri - Goniobranchus splendidus | 2.068 | 0.039 | 1 | Chromodoris kuiteri - Goniobranchus splendidus | 3.154 | 0.002 | 0.126 |
| Discodoris sp - Goniobranchus splendidus | -5.663 | <0.001 | <0.001 | Discodoris sp - Goniobranchus splendidus | -5.923 | <0.001 | <0.001 | Discodoris sp - Goniobranchus splendidus | -5.244 | <0.001 | <0.001 | Discodoris sp - Goniobranchus splendidus | -0.102 | 0.919 | 1 |
| Doriprismatica atromarginata - Goniobranchus splendidus | -0.337 | 0.736 | 1 | Doriprismatica atromarginata - Goniobranchus splendidus | -2.205 | 0.027 | 1 | Doriprismatica atromarginata - Goniobranchus splendidus | -3.716 | 0 | 0.016 | Doriprismatica atromarginata - Goniobranchus splendidus | -1.578 | 0.115 | 1 |
| Glossodoris vespa - Goniobranchus splendidus | -1.118 | 0.264 | 1 | Glossodoris vespa - Goniobranchus splendidus | -0.401 | 0.689 | 1 | Glossodoris vespa - Goniobranchus splendidus | 0.089 | 0.929 | 1 | Glossodoris vespa - Goniobranchus splendidus | 0.345 | 0.73 | 1 |
| Goniobranchus collingwoodi - Goniobranchus splendidus | -6.256 | <0.001 | <0.001 | Goniobranchus collingwoodi - Goniobranchus splendidus | -6.086 | <0.001 | <0.001 | Goniobranchus collingwoodi - Goniobranchus splendidus | -5.147 | <0.001 | <0.001 | Goniobranchus collingwoodi - Goniobranchus splendidus | 0.529 | 0.597 | 1 |
| Aphelodoris varia - Hypselodoris bennetti | 0.483 | 0.629 | 1 | Aphelodoris varia - Hypselodoris bennetti | 1.036 | 0.3 | 1 | Aphelodoris varia - Hypselodoris bennetti | 2.215 | 0.027 | 1 | Aphelodoris varia - Hypselodoris bennetti | 3.845 | <0.001 | 0.009 |
| Chromodoris elisabethina - Hypselodoris bennetti | 2.93 | 0.003 | 0.265 | Chromodoris elisabethina - Hypselodoris bennetti | 2.984 | 0.003 | 0.222 | Chromodoris elisabethina - Hypselodoris bennetti | 3.199 | 0.001 | 0.108 | Chromodoris elisabethina - Hypselodoris bennetti | 3.521 | <0.001 | 0.033 |
| Chromodoris kuiteri - Hypselodoris bennetti | 4.927 | <0.001 | <0.001 | Chromodoris kuiteri - Hypselodoris bennetti | 5.561 | <0.001 | <0.001 | Chromodoris kuiteri - Hypselodoris bennetti | 6.292 | <0.001 | <0.001 | Chromodoris kuiteri - Hypselodoris bennetti | 5.955 | <0.001 | <0.001 |
| Discodoris sp - Hypselodoris bennetti | -1.005 | 0.315 | 1 | Discodoris sp - Hypselodoris bennetti | -0.679 | 0.497 | 1 | Discodoris sp - Hypselodoris bennetti | 0.355 | 0.723 | 1 | Discodoris sp - Hypselodoris bennetti | 3.303 | 0.001 | 0.075 |
| Doriprismatica atromarginata - Hypselodoris bennetti | 3.635 | <0.001 | 0.022 | Doriprismatica atromarginata - Hypselodoris bennetti | 2.824 | 0.005 | 0.37 | Doriprismatica atromarginata - Hypselodoris bennetti | 2.231 | 0.026 | 1 | Doriprismatica atromarginata - Hypselodoris bennetti | 2.549 | 0.011 | 0.842 |
| Glossodoris vespa - Hypselodoris bennetti | 2.631 | 0.009 | 0.664 | Glossodoris vespa - Hypselodoris bennetti | 3.739 | <0.001 | 0.014 | Glossodoris vespa - Hypselodoris bennetti | 4.62 | <0.001 | <0.001 | Glossodoris vespa - Hypselodoris bennetti | 3.66 | <0.001 | 0.02 |
| Goniobranchus collingwoodi - Hypselodoris bennetti | -1.48 | 0.139 | 1 | Goniobranchus collingwoodi - Hypselodoris bennetti | -0.809 | 0.418 | 1 | Goniobranchus collingwoodi - Hypselodoris bennetti | 0.432 | 0.666 | 1 | Goniobranchus collingwoodi - Hypselodoris bennetti | 3.807 | <0.001 | 0.011 |
| Goniobranchus splendidus - Hypselodoris bennetti | 3.846 | <0.001 | 0.009 | Goniobranchus splendidus - Hypselodoris bennetti | 4.429 | <0.001 | 0.001 | Goniobranchus splendidus - Hypselodoris bennetti | 4.964 | <0.001 | <0.001 | Goniobranchus splendidus - Hypselodoris bennetti | 3.693 | <0.001 | 0.017 |
| Aphelodoris varia - Phyllidia elegans | 1.349 | 0.177 | 1 | Aphelodoris varia - Phyllidia elegans | 1.673 | 0.094 | 1 | Aphelodoris varia - Phyllidia elegans | 2.179 | 0.029 | 1 | Aphelodoris varia - Phyllidia elegans | 2.39 | 0.017 | 1 |
| Chromodoris elisabethina - Phyllidia elegans | 3.597 | <0.001 | 0.025 | Chromodoris elisabethina - Phyllidia elegans | 3.465 | 0.001 | 0.041 | Chromodoris elisabethina - Phyllidia elegans | 3.093 | 0.002 | 0.155 | Chromodoris elisabethina - Phyllidia elegans | 2.122 | 0.034 | 1 |
| Chromodoris kuiteri - Phyllidia elegans | 5.44 | <0.001 | <0.001 | Chromodoris kuiteri - Phyllidia elegans | 5.853 | <0.001 | <0.001 | Chromodoris kuiteri - Phyllidia elegans | 5.979 | <0.001 | <0.001 | Chromodoris kuiteri - Phyllidia elegans | 4.436 | <0.001 | 0.001 |
| Discodoris sp - Phyllidia elegans | -0.095 | 0.925 | 1 | Discodoris sp - Phyllidia elegans | 0.036 | 0.971 | 1 | Discodoris sp - Phyllidia elegans | 0.458 | 0.647 | 1 | Discodoris sp - Phyllidia elegans | 2.003 | 0.045 | 1 |
| Doriprismatica atromarginata - Phyllidia elegans | 4.259 | <0.001 | 0.002 | Doriprismatica atromarginata - Phyllidia elegans | 3.325 | 0.001 | 0.069 | Doriprismatica atromarginata - Phyllidia elegans | 2.191 | 0.028 | 1 | Doriprismatica atromarginata - Phyllidia elegans | 1.173 | 0.241 | 1 |
| Glossodoris vespa - Phyllidia elegans | 3.296 | 0.001 | 0.076 | Glossodoris vespa - Phyllidia elegans | 4.156 | <0.001 | 0.003 | Glossodoris vespa - Phyllidia elegans | 4.436 | <0.001 | 0.001 | Glossodoris vespa - Phyllidia elegans | 2.336 | 0.019 | 1 |
| Goniobranchus collingwoodi - Phyllidia elegans | -0.537 | 0.591 | 1 | Goniobranchus collingwoodi - Phyllidia elegans | -0.085 | 0.932 | 1 | Goniobranchus collingwoodi - Phyllidia elegans | 0.53 | 0.596 | 1 | Goniobranchus collingwoodi - Phyllidia elegans | 2.474 | 0.013 | 1 |
| Goniobranchus splendidus - Phyllidia elegans | 4.451 | <0.001 | 0.001 | Goniobranchus splendidus - Phyllidia elegans | 4.802 | <0.001 | <0.001 | Goniobranchus splendidus - Phyllidia elegans | 4.71 | <0.001 | <0.001 | Goniobranchus splendidus - Phyllidia elegans | 2.241 | 0.025 | 1 |
| Hypselodoris bennetti - Phyllidia elegans | 0.778 | 0.437 | 1 | Hypselodoris bennetti - Phyllidia elegans | 0.618 | 0.536 | 1 | Hypselodoris bennetti - Phyllidia elegans | 0.118 | 0.906 | 1 | Hypselodoris bennetti - Phyllidia elegans | -0.994 | 0.32 | 1 |
| Aphelodoris varia - Phyllidia ocellata | -2.954 | 0.003 | 0.244 | Aphelodoris varia - Phyllidia ocellata | -3.257 | 0.001 | 0.088 | Aphelodoris varia - Phyllidia ocellata | -3.218 | 0.001 | 0.101 | Aphelodoris varia - Phyllidia ocellata | 2.594 | 0.009 | 0.741 |
| Chromodoris elisabethina - Phyllidia ocellata | 0.271 | 0.786 | 1 | Chromodoris elisabethina - Phyllidia ocellata | -0.642 | 0.521 | 1 | Chromodoris elisabethina - Phyllidia ocellata | -1.801 | 0.072 | 1 | Chromodoris elisabethina - Phyllidia ocellata | 2.195 | 0.028 | 1 |
| Chromodoris kuiteri - Phyllidia ocellata | 2.876 | 0.004 | 0.314 | Chromodoris kuiteri - Phyllidia ocellata | 2.739 | 0.006 | 0.481 | Chromodoris kuiteri - Phyllidia ocellata | 2.298 | 0.022 | 1 | Chromodoris kuiteri - Phyllidia ocellata | 5.262 | <0.001 | <0.001 |
| Discodoris sp - Phyllidia ocellata | -4.381 | <0.001 | 0.001 | Discodoris sp - Phyllidia ocellata | -4.874 | <0.001 | <0.001 | Discodoris sp - Phyllidia ocellata | -4.905 | <0.001 | <0.001 | Discodoris sp - Phyllidia ocellata | 1.983 | 0.047 | 1 |
| Doriprismatica atromarginata - Phyllidia ocellata | 1.064 | 0.287 | 1 | Doriprismatica atromarginata - Phyllidia ocellata | -1.04 | 0.299 | 1 | Doriprismatica atromarginata - Phyllidia ocellata | -3.336 | 0.001 | 0.066 | Doriprismatica atromarginata - Phyllidia ocellata | 0.893 | 0.372 | 1 |
| Glossodoris vespa - Phyllidia ocellata | 0.092 | 0.927 | 1 | Glossodoris vespa - Phyllidia ocellata | 0.561 | 0.575 | 1 | Glossodoris vespa - Phyllidia ocellata | 0.342 | 0.732 | 1 | Glossodoris vespa - Phyllidia ocellata | 2.422 | 0.015 | 1 |
| Goniobranchus collingwoodi - Phyllidia ocellata | -4.965 | <0.001 | <0.001 | Goniobranchus collingwoodi - Phyllidia ocellata | -5.034 | <0.001 | <0.001 | Goniobranchus collingwoodi - Phyllidia ocellata | -4.81 | <0.001 | <0.001 | Goniobranchus collingwoodi - Phyllidia ocellata | 2.604 | 0.009 | 0.719 |
| Goniobranchus splendidus - Phyllidia ocellata | 1.369 | 0.171 | 1 | Goniobranchus splendidus - Phyllidia ocellata | 1.097 | 0.273 | 1 | Goniobranchus splendidus - Phyllidia ocellata | 0.293 | 0.77 | 1 | Goniobranchus splendidus - Phyllidia ocellata | 2.393 | 0.017 | 1 |
| Hypselodoris bennetti - Phyllidia ocellata | -2.755 | 0.006 | 0.457 | Hypselodoris bennetti - Phyllidia ocellata | -3.538 | <0.001 | 0.031 | Hypselodoris bennetti - Phyllidia ocellata | -4.68 | <0.001 | <0.001 | Hypselodoris bennetti - Phyllidia ocellata | -1.822 | 0.068 | 1 |
| Phyllidia elegans - Phyllidia ocellata | -3.442 | 0.001 | 0.045 | Phyllidia elegans - Phyllidia ocellata | -3.98 | <0.001 | 0.005 | Phyllidia elegans - Phyllidia ocellata | -4.455 | <0.001 | 0.001 | Phyllidia elegans - Phyllidia ocellata | -0.533 | 0.594 | 1 |
| Aphelodoris varia - Phyllidia varicosa | -3.568 | <0.001 | 0.028 | Aphelodoris varia - Phyllidia varicosa | -3.82 | <0.001 | 0.01 | Aphelodoris varia - Phyllidia varicosa | -3.192 | 0.001 | 0.11 | Aphelodoris varia - Phyllidia varicosa | -0.546 | 0.585 | 1 |
| Chromodoris elisabethina - Phyllidia varicosa | -1.234 | 0.217 | 1 | Chromodoris elisabethina - Phyllidia varicosa | -1.932 | 0.053 | 1 | Chromodoris elisabethina - Phyllidia varicosa | -2.185 | 0.029 | 1 | Chromodoris elisabethina - Phyllidia varicosa | -0.763 | 0.445 | 1 |
| Chromodoris kuiteri - Phyllidia varicosa | 0.758 | 0.448 | 1 | Chromodoris kuiteri - Phyllidia varicosa | 0.623 | 0.533 | 1 | Chromodoris kuiteri - Phyllidia varicosa | 0.864 | 0.387 | 1 | Chromodoris kuiteri - Phyllidia varicosa | 1.64 | 0.101 | 1 |
| Discodoris sp - Phyllidia varicosa | -4.68 | <0.001 | <0.001 | Discodoris sp - Phyllidia varicosa | -5.086 | <0.001 | <0.001 | Discodoris sp - Phyllidia varicosa | -4.55 | <0.001 | <0.001 | Discodoris sp - Phyllidia varicosa | -0.735 | 0.462 | 1 |
| Doriprismatica atromarginata - Phyllidia varicosa | -0.727 | 0.467 | 1 | Doriprismatica atromarginata - Phyllidia varicosa | -2.246 | 0.025 | 1 | Doriprismatica atromarginata - Phyllidia varicosa | -3.257 | 0.001 | 0.088 | Doriprismatica atromarginata - Phyllidia varicosa | -1.805 | 0.071 | 1 |
| Glossodoris vespa - Phyllidia varicosa | -1.289 | 0.197 | 1 | Glossodoris vespa - Phyllidia varicosa | -0.966 | 0.334 | 1 | Glossodoris vespa - Phyllidia varicosa | -0.573 | 0.567 | 1 | Glossodoris vespa - Phyllidia varicosa | -0.402 | 0.688 | 1 |
| Goniobranchus collingwoodi - Phyllidia varicosa | -5.122 | <0.001 | <0.001 | Goniobranchus collingwoodi - Phyllidia varicosa | -5.207 | <0.001 | <0.001 | Goniobranchus collingwoodi - Phyllidia varicosa | -4.478 | <0.001 | 0.001 | Goniobranchus collingwoodi - Phyllidia varicosa | -0.264 | 0.792 | 1 |
| Goniobranchus splendidus - Phyllidia varicosa | -0.49 | 0.624 | 1 | Goniobranchus splendidus - Phyllidia varicosa | -0.719 | 0.472 | 1 | Goniobranchus splendidus - Phyllidia varicosa | -0.688 | 0.491 | 1 | Goniobranchus splendidus - Phyllidia varicosa | -0.71 | 0.478 | 1 |
| Hypselodoris bennetti - Phyllidia varicosa | -3.454 | 0.001 | 0.043 | Hypselodoris bennetti - Phyllidia varicosa | -4.109 | <0.001 | 0.003 | Hypselodoris bennetti - Phyllidia varicosa | -4.505 | <0.001 | 0.001 | Hypselodoris bennetti - Phyllidia varicosa | -3.521 | <0.001 | 0.034 |
| Phyllidia elegans - Phyllidia varicosa | -4.015 | <0.001 | 0.005 | Phyllidia elegans - Phyllidia varicosa | -4.485 | <0.001 | 0.001 | Phyllidia elegans - Phyllidia varicosa | -4.386 | <0.001 | 0.001 | Phyllidia elegans - Phyllidia varicosa | -2.397 | 0.017 | 1 |
| Phyllidia ocellata - Phyllidia varicosa | -1.449 | 0.147 | 1 | Phyllidia ocellata - Phyllidia varicosa | -1.484 | 0.138 | 1 | Phyllidia ocellata - Phyllidia varicosa | -0.888 | 0.375 | 1 | Phyllidia ocellata - Phyllidia varicosa | -2.387 | 0.017 | 1 |
| Aphelodoris varia - Phyllidiella pustulosa | 2.318 | 0.02 | 1 | Aphelodoris varia - Phyllidiella pustulosa | 2.336 | 0.019 | 1 | Aphelodoris varia - Phyllidiella pustulosa | 2.408 | 0.016 | 1 | Aphelodoris varia - Phyllidiella pustulosa | 3.863 | <0.001 | 0.009 |
| Chromodoris elisabethina - Phyllidiella pustulosa | 5.189 | <0.001 | <0.001 | Chromodoris elisabethina - Phyllidiella pustulosa | 4.623 | <0.001 | <0.001 | Chromodoris elisabethina - Phyllidiella pustulosa | 3.568 | <0.001 | 0.028 | Chromodoris elisabethina - Phyllidiella pustulosa | 3.457 | 0.001 | 0.043 |
| Chromodoris kuiteri - Phyllidiella pustulosa | 7.406 | <0.001 | <0.001 | Chromodoris kuiteri - Phyllidiella pustulosa | 7.554 | <0.001 | <0.001 | Chromodoris kuiteri - Phyllidiella pustulosa | 7.169 | <0.001 | <0.001 | Chromodoris kuiteri - Phyllidiella pustulosa | 6.3 | <0.001 | <0.001 |
| Discodoris sp - Phyllidiella pustulosa | 0.374 | 0.708 | 1 | Discodoris sp - Phyllidiella pustulosa | 0.175 | 0.861 | 1 | Discodoris sp - Phyllidiella pustulosa | 0.176 | 0.86 | 1 | Discodoris sp - Phyllidiella pustulosa | 3.163 | 0.002 | 0.122 |
| Doriprismatica atromarginata - Phyllidiella pustulosa | 6.201 | <0.001 | <0.001 | Doriprismatica atromarginata - Phyllidiella pustulosa | 4.548 | <0.001 | <0.001 | Doriprismatica atromarginata - Phyllidiella pustulosa | 2.441 | 0.015 | 1 | Doriprismatica atromarginata - Phyllidiella pustulosa | 2.304 | 0.021 | 1 |
| Glossodoris vespa - Phyllidiella pustulosa | 4.621 | <0.001 | <0.001 | Glossodoris vespa - Phyllidiella pustulosa | 5.334 | <0.001 | <0.001 | Glossodoris vespa - Phyllidiella pustulosa | 5.158 | <0.001 | <0.001 | Glossodoris vespa - Phyllidiella pustulosa | 3.58 | <0.001 | 0.027 |
| Goniobranchus collingwoodi - Phyllidiella pustulosa | -0.18 | 0.857 | 1 | Goniobranchus collingwoodi - Phyllidiella pustulosa | 0.023 | 0.981 | 1 | Goniobranchus collingwoodi - Phyllidiella pustulosa | 0.267 | 0.79 | 1 | Goniobranchus collingwoodi - Phyllidiella pustulosa | 3.752 | <0.001 | 0.014 |
| Goniobranchus splendidus - Phyllidiella pustulosa | 6.407 | <0.001 | <0.001 | Goniobranchus splendidus - Phyllidiella pustulosa | 6.456 | <0.001 | <0.001 | Goniobranchus splendidus - Phyllidiella pustulosa | 5.74 | <0.001 | <0.001 | Goniobranchus splendidus - Phyllidiella pustulosa | 3.685 | <0.001 | 0.018 |
| Hypselodoris bennetti - Phyllidiella pustulosa | 1.372 | 0.17 | 1 | Hypselodoris bennetti - Phyllidiella pustulosa | 0.858 | 0.391 | 1 | Hypselodoris bennetti - Phyllidiella pustulosa | -0.211 | 0.833 | 1 | Hypselodoris bennetti - Phyllidiella pustulosa | -0.615 | 0.538 | 1 |
| Phyllidia elegans - Phyllidiella pustulosa | 0.405 | 0.685 | 1 | Phyllidia elegans - Phyllidiella pustulosa | 0.106 | 0.915 | 1 | Phyllidia elegans - Phyllidiella pustulosa | -0.327 | 0.744 | 1 | Phyllidia elegans - Phyllidiella pustulosa | 0.538 | 0.59 | 1 |
| Phyllidia ocellata - Phyllidiella pustulosa | 5.036 | <0.001 | <0.001 | Phyllidia ocellata - Phyllidiella pustulosa | 5.335 | <0.001 | <0.001 | Phyllidia ocellata - Phyllidiella pustulosa | 5.369 | <0.001 | <0.001 | Phyllidia ocellata - Phyllidiella pustulosa | 1.422 | 0.155 | 1 |
| Phyllidia varicosa - Phyllidiella pustulosa | 5.13 | <0.001 | <0.001 | Phyllidia varicosa - Phyllidiella pustulosa | 5.384 | <0.001 | <0.001 | Phyllidia varicosa - Phyllidiella pustulosa | 4.834 | <0.001 | <0.001 | Phyllidia varicosa - Phyllidiella pustulosa | 3.359 | 0.001 | 0.061 |

**Table S4.** Dunn tests at species level for chromatic boldness (Col.CoV). Significant (Bonferroni adjusted p) pairwise comparisons in red.

| **2cm** | | | | **5cm** | | | | **10cm** | | | | **30cm** | | | |
| --- | --- | --- | --- | --- | --- | --- | --- | --- | --- | --- | --- | --- | --- | --- | --- |
| Comparison | Z | P.unadj | P.adj | Comparison | Z | P.unadj | P.adj | Comparison | Z | P.unadj | P.adj | Comparison | Z | P.unadj | P.adj |
| Aphelodoris varia - Chromodoris elisabethina | -3.147 | 0.002 | 0.129 | Aphelodoris varia - Chromodoris elisabethina | -3.465 | 0.001 | 0.041 | Aphelodoris varia - Chromodoris elisabethina | -3.586 | <0.001 | 0.026 | Aphelodoris varia - Chromodoris elisabethina | -2.795 | 0.005 | 0.405 |
| Aphelodoris varia - Chromodoris kuiteri | -4.663 | <0.001 | <0.001 | Aphelodoris varia - Chromodoris kuiteri | -4.808 | <0.001 | <0.001 | Aphelodoris varia - Chromodoris kuiteri | -4.999 | <0.001 | <0.001 | Aphelodoris varia - Chromodoris kuiteri | -4.826 | <0.001 | <0.001 |
| Chromodoris elisabethina - Chromodoris kuiteri | -1.649 | 0.099 | 1 | Chromodoris elisabethina - Chromodoris kuiteri | -1.498 | 0.134 | 1 | Chromodoris elisabethina - Chromodoris kuiteri | -1.574 | 0.116 | 1 | Chromodoris elisabethina - Chromodoris kuiteri | -2.129 | 0.033 | 1 |
| Aphelodoris varia - Discodoris sp | 1.36 | 0.174 | 1 | Aphelodoris varia - Discodoris sp | 1.255 | 0.209 | 1 | Aphelodoris varia - Discodoris sp | 0.879 | 0.38 | 1 | Aphelodoris varia - Discodoris sp | -0.821 | 0.412 | 1 |
| Chromodoris elisabethina - Discodoris sp | 4.105 | <0.001 | 0.003 | Chromodoris elisabethina - Discodoris sp | 4.284 | <0.001 | 0.001 | Chromodoris elisabethina - Discodoris sp | 4.025 | <0.001 | 0.004 | Chromodoris elisabethina - Discodoris sp | 1.671 | 0.095 | 1 |
| Chromodoris kuiteri - Discodoris sp | 5.436 | <0.001 | <0.001 | Chromodoris kuiteri - Discodoris sp | 5.468 | <0.001 | <0.001 | Chromodoris kuiteri - Discodoris sp | 5.29 | <0.001 | <0.001 | Chromodoris kuiteri - Discodoris sp | 3.556 | <0.001 | 0.029 |
| Aphelodoris varia - Doriprismatica atromarginata | -1.381 | 0.167 | 1 | Aphelodoris varia - Doriprismatica atromarginata | -0.344 | 0.731 | 1 | Aphelodoris varia - Doriprismatica atromarginata | 1.434 | 0.152 | 1 | Aphelodoris varia - Doriprismatica atromarginata | 0.084 | 0.933 | 1 |
| Chromodoris elisabethina - Doriprismatica atromarginata | 1.9 | 0.057 | 1 | Chromodoris elisabethina - Doriprismatica atromarginata | 3.227 | 0.001 | 0.098 | Chromodoris elisabethina - Doriprismatica atromarginata | 5.065 | <0.001 | <0.001 | Chromodoris elisabethina - Doriprismatica atromarginata | 2.951 | 0.003 | 0.247 |
| Chromodoris kuiteri - Doriprismatica atromarginata | 3.523 | <0.001 | 0.033 | Chromodoris kuiteri - Doriprismatica atromarginata | 4.611 | <0.001 | <0.001 | Chromodoris kuiteri - Doriprismatica atromarginata | 6.418 | <0.001 | <0.001 | Chromodoris kuiteri - Doriprismatica atromarginata | 5.017 | <0.001 | <0.001 |
| Discodoris sp - Doriprismatica atromarginata | -2.593 | 0.01 | 0.742 | Discodoris sp - Doriprismatica atromarginata | -1.582 | 0.114 | 1 | Discodoris sp - Doriprismatica atromarginata | 0.351 | 0.726 | 1 | Discodoris sp - Doriprismatica atromarginata | 0.912 | 0.362 | 1 |
| Aphelodoris varia - Glossodoris vespa | -5.353 | <0.001 | <0.001 | Aphelodoris varia - Glossodoris vespa | -5.229 | <0.001 | <0.001 | Aphelodoris varia - Glossodoris vespa | -5.023 | <0.001 | <0.001 | Aphelodoris varia - Glossodoris vespa | -4.832 | <0.001 | <0.001 |
| Chromodoris elisabethina - Glossodoris vespa | -2.43 | 0.015 | 1 | Chromodoris elisabethina - Glossodoris vespa | -2.028 | 0.043 | 1 | Chromodoris elisabethina - Glossodoris vespa | -1.721 | 0.085 | 1 | Chromodoris elisabethina - Glossodoris vespa | -2.234 | 0.025 | 1 |
| Chromodoris kuiteri - Glossodoris vespa | -0.801 | 0.423 | 1 | Chromodoris kuiteri - Glossodoris vespa | -0.556 | 0.578 | 1 | Chromodoris kuiteri - Glossodoris vespa | -0.193 | 0.847 | 1 | Chromodoris kuiteri - Glossodoris vespa | -0.171 | 0.864 | 1 |
| Discodoris sp - Glossodoris vespa | -6.051 | <0.001 | <0.001 | Discodoris sp - Glossodoris vespa | -5.844 | <0.001 | <0.001 | Discodoris sp - Glossodoris vespa | -5.319 | <0.001 | <0.001 | Discodoris sp - Glossodoris vespa | -3.616 | <0.001 | 0.023 |
| Doriprismatica atromarginata - Glossodoris vespa | -4.268 | <0.001 | 0.002 | Doriprismatica atromarginata - Glossodoris vespa | -5.044 | <0.001 | <0.001 | Doriprismatica atromarginata - Glossodoris vespa | -6.383 | <0.001 | <0.001 | Doriprismatica atromarginata - Glossodoris vespa | -5.012 | <0.001 | <0.001 |
| Aphelodoris varia - Goniobranchus collingwoodi | 2.565 | 0.01 | 0.805 | Aphelodoris varia - Goniobranchus collingwoodi | 2.541 | 0.011 | 0.863 | Aphelodoris varia - Goniobranchus collingwoodi | 2.164 | 0.03 | 1 | Aphelodoris varia - Goniobranchus collingwoodi | -1.282 | 0.2 | 1 |
| Chromodoris elisabethina - Goniobranchus collingwoodi | 5.278 | <0.001 | <0.001 | Chromodoris elisabethina - Goniobranchus collingwoodi | 5.536 | <0.001 | <0.001 | Chromodoris elisabethina - Goniobranchus collingwoodi | 5.276 | <0.001 | <0.001 | Chromodoris elisabethina - Goniobranchus collingwoodi | 1.222 | 0.222 | 1 |
| Chromodoris kuiteri - Goniobranchus collingwoodi | 6.556 | <0.001 | <0.001 | Chromodoris kuiteri - Goniobranchus collingwoodi | 6.663 | <0.001 | <0.001 | Chromodoris kuiteri - Goniobranchus collingwoodi | 6.485 | <0.001 | <0.001 | Chromodoris kuiteri - Goniobranchus collingwoodi | 3.127 | 0.002 | 0.138 |
| Discodoris sp - Goniobranchus collingwoodi | 1.086 | 0.277 | 1 | Discodoris sp - Goniobranchus collingwoodi | 1.159 | 0.247 | 1 | Discodoris sp - Goniobranchus collingwoodi | 1.159 | 0.247 | 1 | Discodoris sp - Goniobranchus collingwoodi | -0.416 | 0.677 | 1 |
| Doriprismatica atromarginata - Goniobranchus collingwoodi | 3.825 | <0.001 | 0.01 | Doriprismatica atromarginata - Goniobranchus collingwoodi | 2.896 | 0.004 | 0.294 | Doriprismatica atromarginata - Goniobranchus collingwoodi | 0.963 | 0.336 | 1 | Doriprismatica atromarginata - Goniobranchus collingwoodi | -1.384 | 0.166 | 1 |
| Glossodoris vespa - Goniobranchus collingwoodi | 7.137 | <0.001 | <0.001 | Glossodoris vespa - Goniobranchus collingwoodi | 7.003 | <0.001 | <0.001 | Glossodoris vespa - Goniobranchus collingwoodi | 6.478 | <0.001 | <0.001 | Glossodoris vespa - Goniobranchus collingwoodi | 3.2 | 0.001 | 0.107 |
| Aphelodoris varia - Goniobranchus splendidus | -3.38 | 0.001 | 0.056 | Aphelodoris varia - Goniobranchus splendidus | -2.837 | 0.005 | 0.356 | Aphelodoris varia - Goniobranchus splendidus | -1.729 | 0.084 | 1 | Aphelodoris varia - Goniobranchus splendidus | -1.186 | 0.236 | 1 |
| Chromodoris elisabethina - Goniobranchus splendidus | -0.087 | 0.931 | 1 | Chromodoris elisabethina - Goniobranchus splendidus | 0.759 | 0.448 | 1 | Chromodoris elisabethina - Goniobranchus splendidus | 1.95 | 0.051 | 1 | Chromodoris elisabethina - Goniobranchus splendidus | 1.677 | 0.094 | 1 |
| Chromodoris kuiteri - Goniobranchus splendidus | 1.629 | 0.103 | 1 | Chromodoris kuiteri - Goniobranchus splendidus | 2.27 | 0.023 | 1 | Chromodoris kuiteri - Goniobranchus splendidus | 3.47 | 0.001 | 0.041 | Chromodoris kuiteri - Goniobranchus splendidus | 3.789 | <0.001 | 0.012 |
| Discodoris sp - Goniobranchus splendidus | -4.328 | <0.001 | 0.001 | Discodoris sp - Goniobranchus splendidus | -3.747 | <0.001 | 0.014 | Discodoris sp - Goniobranchus splendidus | -2.398 | 0.016 | 1 | Discodoris sp - Goniobranchus splendidus | -0.21 | 0.833 | 1 |
| Doriprismatica atromarginata - Goniobranchus splendidus | -2.084 | 0.037 | 1 | Doriprismatica atromarginata - Goniobranchus splendidus | -2.573 | 0.01 | 0.786 | Doriprismatica atromarginata - Goniobranchus splendidus | -3.229 | 0.001 | 0.097 | Doriprismatica atromarginata - Goniobranchus splendidus | -1.306 | 0.192 | 1 |
| Glossodoris vespa - Goniobranchus splendidus | 2.437 | 0.015 | 1 | Glossodoris vespa - Goniobranchus splendidus | 2.787 | 0.005 | 0.415 | Glossodoris vespa - Goniobranchus splendidus | 3.549 | <0.001 | 0.03 | Glossodoris vespa - Goniobranchus splendidus | 3.832 | <0.001 | 0.01 |
| Goniobranchus collingwoodi - Goniobranchus splendidus | -5.543 | <0.001 | <0.001 | Goniobranchus collingwoodi - Goniobranchus splendidus | -5.042 | <0.001 | <0.001 | Goniobranchus collingwoodi - Goniobranchus splendidus | -3.694 | 0 | 0.017 | Goniobranchus collingwoodi - Goniobranchus splendidus | 0.255 | 0.799 | 1 |
| Aphelodoris varia - Hypselodoris bennetti | -3.564 | <0.001 | 0.029 | Aphelodoris varia - Hypselodoris bennetti | -3.231 | 0.001 | 0.096 | Aphelodoris varia - Hypselodoris bennetti | -0.042 | 0.967 | 1 | Aphelodoris varia - Hypselodoris bennetti | 1.207 | 0.227 | 1 |
| Chromodoris elisabethina - Hypselodoris bennetti | -1.044 | 0.297 | 1 | Chromodoris elisabethina - Hypselodoris bennetti | -0.471 | 0.638 | 1 | Chromodoris elisabethina - Hypselodoris bennetti | 2.748 | 0.006 | 0.468 | Chromodoris elisabethina - Hypselodoris bennetti | 3.356 | 0.001 | 0.062 |
| Chromodoris kuiteri - Hypselodoris bennetti | 0.343 | 0.731 | 1 | Chromodoris kuiteri - Hypselodoris bennetti | 0.772 | 0.44 | 1 | Chromodoris kuiteri - Hypselodoris bennetti | 3.937 | <0.001 | 0.006 | Chromodoris kuiteri - Hypselodoris bennetti | 4.979 | <0.001 | <0.001 |
| Discodoris sp - Hypselodoris bennetti | -4.382 | <0.001 | 0.001 | Discodoris sp - Hypselodoris bennetti | -3.991 | <0.001 | 0.005 | Discodoris sp - Hypselodoris bennetti | -0.747 | 0.455 | 1 | Discodoris sp - Hypselodoris bennetti | 1.774 | 0.076 | 1 |
| Doriprismatica atromarginata - Hypselodoris bennetti | -2.577 | 0.01 | 0.778 | Doriprismatica atromarginata - Hypselodoris bennetti | -3.025 | 0.002 | 0.194 | Doriprismatica atromarginata - Hypselodoris bennetti | -1.129 | 0.259 | 1 | Doriprismatica atromarginata - Hypselodoris bennetti | 1.164 | 0.245 | 1 |
| Glossodoris vespa - Hypselodoris bennetti | 1.03 | 0.303 | 1 | Glossodoris vespa - Hypselodoris bennetti | 1.236 | 0.216 | 1 | Glossodoris vespa - Hypselodoris bennetti | 4.011 | <0.001 | 0.005 | Glossodoris vespa - Hypselodoris bennetti | 5.009 | <0.001 | <0.001 |
| Goniobranchus collingwoodi - Hypselodoris bennetti | -5.353 | <0.001 | <0.001 | Goniobranchus collingwoodi - Hypselodoris bennetti | -5.027 | <0.001 | <0.001 | Goniobranchus collingwoodi - Hypselodoris bennetti | -1.783 | 0.075 | 1 | Goniobranchus collingwoodi - Hypselodoris bennetti | 2.147 | 0.032 | 1 |
| Goniobranchus splendidus - Hypselodoris bennetti | -1.003 | 0.316 | 1 | Goniobranchus splendidus - Hypselodoris bennetti | -1.084 | 0.278 | 1 | Goniobranchus splendidus - Hypselodoris bennetti | 1.279 | 0.201 | 1 | Goniobranchus splendidus - Hypselodoris bennetti | 2.12 | 0.034 | 1 |
| Aphelodoris varia - Phyllidia elegans | 2.149 | 0.032 | 1 | Aphelodoris varia - Phyllidia elegans | 2.295 | 0.022 | 1 | Aphelodoris varia - Phyllidia elegans | 2.084 | 0.037 | 1 | Aphelodoris varia - Phyllidia elegans | 0.968 | 0.333 | 1 |
| Chromodoris elisabethina - Phyllidia elegans | 4.375 | <0.001 | 0.001 | Chromodoris elisabethina - Phyllidia elegans | 4.747 | <0.001 | <0.001 | Chromodoris elisabethina - Phyllidia elegans | 4.626 | <0.001 | <0.001 | Chromodoris elisabethina - Phyllidia elegans | 2.961 | 0.003 | 0.239 |
| Chromodoris kuiteri - Phyllidia elegans | 5.494 | <0.001 | <0.001 | Chromodoris kuiteri - Phyllidia elegans | 5.74 | <0.001 | <0.001 | Chromodoris kuiteri - Phyllidia elegans | 5.681 | <0.001 | <0.001 | Chromodoris kuiteri - Phyllidia elegans | 4.49 | <0.001 | 0.001 |
| Discodoris sp - Phyllidia elegans | 0.982 | 0.326 | 1 | Discodoris sp - Phyllidia elegans | 1.196 | 0.232 | 1 | Discodoris sp - Phyllidia elegans | 1.283 | 0.2 | 1 | Discodoris sp - Phyllidia elegans | 1.519 | 0.129 | 1 |
| Doriprismatica atromarginata - Phyllidia elegans | 3.142 | 0.002 | 0.131 | Doriprismatica atromarginata - Phyllidia elegans | 2.567 | 0.01 | 0.8 | Doriprismatica atromarginata - Phyllidia elegans | 1.114 | 0.265 | 1 | Doriprismatica atromarginata - Phyllidia elegans | 0.923 | 0.356 | 1 |
| Glossodoris vespa - Phyllidia elegans | 6.029 | <0.001 | <0.001 | Glossodoris vespa - Phyllidia elegans | 6.071 | <0.001 | <0.001 | Glossodoris vespa - Phyllidia elegans | 5.719 | <0.001 | <0.001 | Glossodoris vespa - Phyllidia elegans | 4.535 | <0.001 | <0.001 |
| Goniobranchus collingwoodi - Phyllidia elegans | 0.076 | 0.939 | 1 | Goniobranchus collingwoodi - Phyllidia elegans | 0.23 | 0.818 | 1 | Goniobranchus collingwoodi - Phyllidia elegans | 0.316 | 0.752 | 1 | Goniobranchus collingwoodi - Phyllidia elegans | 1.866 | 0.062 | 1 |
| Goniobranchus splendidus - Phyllidia elegans | 4.538 | <0.001 | <0.001 | Goniobranchus splendidus - Phyllidia elegans | 4.302 | <0.001 | 0.001 | Goniobranchus splendidus - Phyllidia elegans | 3.311 | 0.001 | 0.073 | Goniobranchus splendidus - Phyllidia elegans | 1.807 | 0.071 | 1 |
| Hypselodoris bennetti - Phyllidia elegans | 4.678 | <0.001 | <0.001 | Hypselodoris bennetti - Phyllidia elegans | 4.539 | <0.001 | <0.001 | Hypselodoris bennetti - Phyllidia elegans | 1.827 | 0.068 | 1 | Hypselodoris bennetti - Phyllidia elegans | -0.125 | 0.901 | 1 |
| Aphelodoris varia - Phyllidia ocellata | 0.508 | 0.611 | 1 | Aphelodoris varia - Phyllidia ocellata | 0.251 | 0.802 | 1 | Aphelodoris varia - Phyllidia ocellata | -0.63 | 0.529 | 1 | Aphelodoris varia - Phyllidia ocellata | -0.75 | 0.453 | 1 |
| Chromodoris elisabethina - Phyllidia ocellata | 3.606 | <0.001 | 0.024 | Chromodoris elisabethina - Phyllidia ocellata | 3.673 | <0.001 | 0.019 | Chromodoris elisabethina - Phyllidia ocellata | 2.941 | 0.003 | 0.255 | Chromodoris elisabethina - Phyllidia ocellata | 2.041 | 0.041 | 1 |
| Chromodoris kuiteri - Phyllidia ocellata | 5.085 | <0.001 | <0.001 | Chromodoris kuiteri - Phyllidia ocellata | 4.994 | <0.001 | <0.001 | Chromodoris kuiteri - Phyllidia ocellata | 4.38 | <0.001 | 0.001 | Chromodoris kuiteri - Phyllidia ocellata | 4.099 | <0.001 | 0.003 |
| Discodoris sp - Phyllidia ocellata | -0.902 | 0.367 | 1 | Discodoris sp - Phyllidia ocellata | -1.024 | 0.306 | 1 | Discodoris sp - Phyllidia ocellata | -1.425 | 0.154 | 1 | Discodoris sp - Phyllidia ocellata | 0.154 | 0.877 | 1 |
| Doriprismatica atromarginata - Phyllidia ocellata | 1.888 | 0.059 | 1 | Doriprismatica atromarginata - Phyllidia ocellata | 0.598 | 0.55 | 1 | Doriprismatica atromarginata - Phyllidia ocellata | -2.065 | 0.039 | 1 | Doriprismatica atromarginata - Phyllidia ocellata | -0.855 | 0.393 | 1 |
| Glossodoris vespa - Phyllidia ocellata | 5.756 | <0.001 | <0.001 | Glossodoris vespa - Phyllidia ocellata | 5.406 | <0.001 | <0.001 | Glossodoris vespa - Phyllidia ocellata | 4.427 | <0.001 | 0.001 | Glossodoris vespa - Phyllidia ocellata | 4.133 | <0.001 | 0.003 |
| Goniobranchus collingwoodi - Phyllidia ocellata | -2.097 | 0.036 | 1 | Goniobranchus collingwoodi - Phyllidia ocellata | -2.299 | 0.022 | 1 | Goniobranchus collingwoodi - Phyllidia ocellata | -2.7 | 0.007 | 0.541 | Goniobranchus collingwoodi - Phyllidia ocellata | 0.612 | 0.541 | 1 |
| Goniobranchus splendidus - Phyllidia ocellata | 3.857 | <0.001 | 0.009 | Goniobranchus splendidus - Phyllidia ocellata | 3.059 | 0.002 | 0.173 | Goniobranchus splendidus - Phyllidia ocellata | 1.074 | 0.283 | 1 | Goniobranchus splendidus - Phyllidia ocellata | 0.415 | 0.678 | 1 |
| Hypselodoris bennetti - Phyllidia ocellata | 3.933 | <0.001 | 0.007 | Hypselodoris bennetti - Phyllidia ocellata | 3.404 | 0.001 | 0.052 | Hypselodoris bennetti - Phyllidia ocellata | -0.444 | 0.657 | 1 | Hypselodoris bennetti - Phyllidia ocellata | -1.777 | 0.075 | 1 |
| Phyllidia elegans - Phyllidia ocellata | -1.777 | 0.076 | 1 | Phyllidia elegans - Phyllidia ocellata | -2.104 | 0.035 | 1 | Phyllidia elegans - Phyllidia ocellata | -2.52 | 0.012 | 0.914 | Phyllidia elegans - Phyllidia ocellata | -1.496 | 0.135 | 1 |
| Aphelodoris varia - Phyllidia varicosa | -1.166 | 0.244 | 1 | Aphelodoris varia - Phyllidia varicosa | -1.986 | 0.047 | 1 | Aphelodoris varia - Phyllidia varicosa | -2.426 | 0.015 | 1 | Aphelodoris varia - Phyllidia varicosa | -1.767 | 0.077 | 1 |
| Chromodoris elisabethina - Phyllidia varicosa | 1.117 | 0.264 | 1 | Chromodoris elisabethina - Phyllidia varicosa | 0.541 | 0.589 | 1 | Chromodoris elisabethina - Phyllidia varicosa | 0.195 | 0.845 | 1 | Chromodoris elisabethina - Phyllidia varicosa | 0.274 | 0.784 | 1 |
| Chromodoris kuiteri - Phyllidia varicosa | 2.337 | 0.019 | 1 | Chromodoris kuiteri - Phyllidia varicosa | 1.664 | 0.096 | 1 | Chromodoris kuiteri - Phyllidia varicosa | 1.387 | 0.166 | 1 | Chromodoris kuiteri - Phyllidia varicosa | 1.886 | 0.059 | 1 |
| Discodoris sp - Phyllidia varicosa | -2.11 | 0.035 | 1 | Discodoris sp - Phyllidia varicosa | -2.795 | 0.005 | 0.405 | Discodoris sp - Phyllidia varicosa | -2.923 | 0.003 | 0.271 | Discodoris sp - Phyllidia varicosa | -1.031 | 0.303 | 1 |
| Doriprismatica atromarginata - Phyllidia varicosa | -0.22 | 0.826 | 1 | Doriprismatica atromarginata - Phyllidia varicosa | -1.774 | 0.076 | 1 | Doriprismatica atromarginata - Phyllidia varicosa | -3.459 | 0.001 | 0.042 | Doriprismatica atromarginata - Phyllidia varicosa | -1.851 | 0.064 | 1 |
| Glossodoris vespa - Phyllidia varicosa | 2.937 | 0.003 | 0.258 | Glossodoris vespa - Phyllidia varicosa | 2.079 | 0.038 | 1 | Glossodoris vespa - Phyllidia varicosa | 1.514 | 0.13 | 1 | Glossodoris vespa - Phyllidia varicosa | 1.985 | 0.047 | 1 |
| Goniobranchus collingwoodi - Phyllidia varicosa | -3.016 | 0.003 | 0.2 | Goniobranchus collingwoodi - Phyllidia varicosa | -3.762 | 0 | 0.013 | Goniobranchus collingwoodi - Phyllidia varicosa | -3.889 | <0.001 | 0.008 | Goniobranchus collingwoodi - Phyllidia varicosa | -0.684 | 0.494 | 1 |
| Goniobranchus splendidus - Phyllidia varicosa | 1.206 | 0.228 | 1 | Goniobranchus splendidus - Phyllidia varicosa | 0 | 1 | 1 | Goniobranchus splendidus - Phyllidia varicosa | -1.222 | 0.222 | 1 | Goniobranchus splendidus - Phyllidia varicosa | -0.942 | 0.346 | 1 |
| Hypselodoris bennetti - Phyllidia varicosa | 1.824 | 0.068 | 1 | Hypselodoris bennetti - Phyllidia varicosa | 0.855 | 0.392 | 1 | Hypselodoris bennetti - Phyllidia varicosa | -2.055 | 0.04 | 1 | Hypselodoris bennetti - Phyllidia varicosa | -2.479 | 0.013 | 1 |
| Phyllidia elegans - Phyllidia varicosa | -2.707 | 0.007 | 0.529 | Phyllidia elegans - Phyllidia varicosa | -3.495 | <0.001 | 0.037 | Phyllidia elegans - Phyllidia varicosa | -3.682 | <0.001 | 0.018 | Phyllidia elegans - Phyllidia varicosa | -2.233 | 0.026 | 1 |
| Phyllidia ocellata - Phyllidia varicosa | -1.521 | 0.128 | 1 | Phyllidia ocellata - Phyllidia varicosa | -2.153 | 0.031 | 1 | Phyllidia ocellata - Phyllidia varicosa | -1.965 | 0.049 | 1 | Phyllidia ocellata - Phyllidia varicosa | -1.224 | 0.221 | 1 |
| Aphelodoris varia - Phyllidiella pustulosa | 3.387 | 0.001 | 0.055 | Aphelodoris varia - Phyllidiella pustulosa | 3.427 | 0.001 | 0.048 | Aphelodoris varia - Phyllidiella pustulosa | 3.098 | 0.002 | 0.152 | Aphelodoris varia - Phyllidiella pustulosa | 2.079 | 0.038 | 1 |
| Chromodoris elisabethina - Phyllidiella pustulosa | 6.215 | <0.001 | <0.001 | Chromodoris elisabethina - Phyllidiella pustulosa | 6.55 | <0.001 | <0.001 | Chromodoris elisabethina - Phyllidiella pustulosa | 6.343 | <0.001 | <0.001 | Chromodoris elisabethina - Phyllidiella pustulosa | 4.617 | <0.001 | <0.001 |
| Chromodoris kuiteri - Phyllidiella pustulosa | 7.493 | <0.001 | <0.001 | Chromodoris kuiteri - Phyllidiella pustulosa | 7.666 | <0.001 | <0.001 | Chromodoris kuiteri - Phyllidiella pustulosa | 7.542 | <0.001 | <0.001 | Chromodoris kuiteri - Phyllidiella pustulosa | 6.44 | <0.001 | <0.001 |
| Discodoris sp - Phyllidiella pustulosa | 1.741 | 0.082 | 1 | Discodoris sp - Phyllidiella pustulosa | 1.875 | 0.061 | 1 | Discodoris sp - Phyllidiella pustulosa | 1.936 | 0.053 | 1 | Discodoris sp - Phyllidiella pustulosa | 2.626 | 0.009 | 0.673 |
| Doriprismatica atromarginata - Phyllidiella pustulosa | 4.744 | <0.001 | <0.001 | Doriprismatica atromarginata - Phyllidiella pustulosa | 3.829 | <0.001 | 0.01 | Doriprismatica atromarginata - Phyllidiella pustulosa | 1.852 | 0.064 | 1 | Doriprismatica atromarginata - Phyllidiella pustulosa | 2.052 | 0.04 | 1 |
| Glossodoris vespa - Phyllidiella pustulosa | 8.061 | <0.001 | <0.001 | Glossodoris vespa - Phyllidiella pustulosa | 7.979 | <0.001 | <0.001 | Glossodoris vespa - Phyllidiella pustulosa | 7.491 | <0.001 | <0.001 | Glossodoris vespa - Phyllidiella pustulosa | 6.403 | <0.001 | <0.001 |
| Goniobranchus collingwoodi - Phyllidiella pustulosa | 0.606 | 0.544 | 1 | Goniobranchus collingwoodi - Phyllidiella pustulosa | 0.665 | 0.506 | 1 | Goniobranchus collingwoodi - Phyllidiella pustulosa | 0.725 | 0.468 | 1 | Goniobranchus collingwoodi - Phyllidiella pustulosa | 3.061 | 0.002 | 0.172 |
| Goniobranchus splendidus - Phyllidiella pustulosa | 6.542 | <0.001 | <0.001 | Goniobranchus splendidus - Phyllidiella pustulosa | 6.08 | <0.001 | <0.001 | Goniobranchus splendidus - Phyllidiella pustulosa | 4.723 | <0.001 | <0.001 | Goniobranchus splendidus - Phyllidiella pustulosa | 3.193 | 0.001 | 0.11 |
| Hypselodoris bennetti - Phyllidiella pustulosa | 6.079 | <0.001 | <0.001 | Hypselodoris bennetti - Phyllidiella pustulosa | 5.793 | <0.001 | <0.001 | Hypselodoris bennetti - Phyllidiella pustulosa | 2.489 | 0.013 | 1 | Hypselodoris bennetti - Phyllidiella pustulosa | 0.491 | 0.623 | 1 |
| Phyllidia elegans - Phyllidiella pustulosa | 0.42 | 0.674 | 1 | Phyllidia elegans - Phyllidiella pustulosa | 0.31 | 0.756 | 1 | Phyllidia elegans - Phyllidiella pustulosa | 0.271 | 0.786 | 1 | Phyllidia elegans - Phyllidiella pustulosa | 0.595 | 0.552 | 1 |
| Phyllidia ocellata - Phyllidiella pustulosa | 2.885 | 0.004 | 0.305 | Phyllidia ocellata - Phyllidiella pustulosa | 3.163 | 0.002 | 0.122 | Phyllidia ocellata - Phyllidiella pustulosa | 3.653 | <0.001 | 0.02 | Phyllidia ocellata - Phyllidiella pustulosa | 2.755 | 0.006 | 0.458 |
| Phyllidia varicosa - Phyllidiella pustulosa | 3.606 | <0.001 | 0.024 | Phyllidia varicosa - Phyllidiella pustulosa | 4.423 | <0.001 | 0.001 | Phyllidia varicosa - Phyllidiella pustulosa | 4.604 | <0.001 | <0.001 | Phyllidia varicosa - Phyllidiella pustulosa | 3.223 | 0.001 | 0.099 |
